# Coupled enzyme discovery, evolution and synthetic yeast chassis adaptation for microbial biopolymer valorisation

**DOI:** 10.64898/2026.09.28.754783

**Authors:** María del Carmen Sánchez Olmos, Jacob Gonzalez Isa, Nicole Paczia, María Mercedes Zambrano, Jaime Huertas Cepas, Eva García-Ruiz, Valerie DeAnda, Daniel Schindler

## Abstract

The valorisation of biological polymers requires microbial systems that can both access recalcitrant substrates and convert the resulting carbon into useful products. Although microbial genome and metagenome resources provide an expanding reservoir of candidate depolymerizing and modifying enzymes, most discovery workflows remain disconnected from enzyme optimisation and host adaptation. Here we present a coupled sequence-based enzyme discovery, enzyme evolution and synthetic yeast chassis adaptation strategy for microbial biopolymer valorisation. Focusing on laccases for the depolymerisation of lignin as a proof of concept, we combine sequence data mining for enzyme discovery, modular yeast surface display for functional screening, directed evolution for enzyme optimisation and synthetic yeast genome diversification for chassis improvement. In our study, surface display enabled functional benchmarking and recovery of improved laccase variants and synthetic-genome-enabled diversification provided a route to explore host configurations that influence display and enzyme performance. By integrating enzyme-level and chassis-level optimisation, this framework addresses a central bottleneck in converting microbial biodiversity by computational sequence repository mining into deployable biomanufacturing systems. Our results establish laccases as tractable entry points for oxidative biopolymer conversion and provide a generalizable platform for engineering yeast systems for sustainable carbon valorisation.

## Introduction

Biological polymers represent one of the largest reservoirs of renewable carbon on Earth; lignocellulosic biomass alone accounts for approximately 430 gigatonnes globally (Bar-On et al., 2018; Boerjan et al., 2003), yet lignocellulosic and other biological polymers remain underused as feedstocks for microbial biotechnology (Ragauskas et al., 2014). Plant biomass and other heterogeneous waste streams contain chemically complex polymers that are abundant, recalcitrant and difficult to convert selectively into defined products (Himmel et al., 2007; Petridis & Smith, 2018). Established industrial valorisation routes largely rely on physical, chemical and thermochemical processing (Ragaert et al., 2017; Saad & Gonçalves, 2024). However, sustainable unlocking of these resources requires biological systems that can depolymerise and transform complex substrates under mild conditions. Microorganisms have evolved diverse strategies to deconstruct and metabolise such materials, making microbial biodiversity a powerful source of catalytic functions for sustainable biomanufacturing (Weiland et al., 2022). However, translating this natural diversity into engineered microbial platforms is still a significant challenge, because enzyme discovery, enzyme optimisation and host adaptation are usually carried out as separate steps. Despite advances in computational enzyme discovery, linking these efforts with experimental validation remains a major bottleneck.

The rapid expansion of genome and metagenome sequence space has transformed the search for enzymes involved in polymer deconstruction (e.g., (Lombard et al., 2014)). Data mining can now be used to identify candidate biocatalysts from organisms that are difficult or impossible to cultivate (Robinson et al., 2021). Meanwhile, DNA synthesis and assembly technologies enable the physical generation of almost any DNA sequence, greatly expanding the range of accessible microbial functions (Hughes & Ellington, 2017; Tan et al., 2025). Sequence-based discovery has been particularly important, yet sequence information alone rarely predicts whether a candidate enzyme will be functional. Although artificial intelligence (AI) can now extract and predict an almost unlimited number of sequences (Ferruz et al., 2022; Madani et al., 2023; Rives et al., 2021; Szymczak et al., 2025), including whole genomes (Brixi et al., 2026), the number of sequences that can be experimentally validated is limited. Often, the majority of predicted sequences are not functional; nevertheless, AI-based sequence prediction can generate synthetic bacteriophage genomes, with approximately 5% of selected sequences being experimentally validated as functional (King et al., 2026). This indicates, despite the rapid progression of computational tools, that functional validation must still be performed experimentally. Consequently, discovery pipelines must be coupled with experimental benchmarking and engineering strategies that can convert sequence diversity into improved, selectable phenotypes, enabling the delivery of products to market (Barber-Zucker et al., 2022).

Laccases (EC 1.10.3.2) are attractive model enzymes for developing such workflows because they catalyse the oxidation of a broad range of phenolic and non-phenolic substrates using molecular oxygen as the terminal electron acceptor (Claus, 2004). Through this chemistry, laccases contribute to the transformation of lignin-derived compounds and other aromatic or polymeric substrates, and have, for example, been explored for applications in biomass processing and bioremediation (Aza & Camarero, 2023; Margot et al., 2015; Weng et al., 2021). Their broad substrate scope makes them promising entry points for biopolymer valorisation, but also creates a need for systematic enzyme selection and optimisation. In addition, choosing an appropriate microbial host and engineering strategy is required to ensure the optimal expression and translation of a given gene or sequence library (Schutz et al., 2023).

Directed evolution provides a route to overcome some limitations by generating and selecting enzyme variants with improved or altered properties (Currin et al., 2015; Garcia-Ruiz et al., 2012; Garcia-Ruiz et al., 2014; Kan et al., 2016). For enzymes acting on complex or heterogeneous substrates, however, the effectiveness of directed evolution depends on the ability to screen large libraries while preserving a reliable link between genotype and phenotype. Yeast surface display offers a powerful solution to this challenge by presenting enzyme variants on the cell surface (Boder & Wittrup, 1997), which can accelerate the benchmarking and evolution of enzymes by avoiding purification, enabling modular library construction and supporting high-throughput selection strategies (Cherf & Cochran, 2015; Li et al., 2024). For laccases and related oxidative enzymes, such systems could provide a direct interface between enzyme discovery, functional screening and variant recovery.

Enzyme optimisation alone, however, is insufficient to establish microbial platforms for biopolymer upcycling. The host chassis must also express the enzyme, tolerate the substrate and by-products, and metabolise depolymerisation products into desired outputs. These requirements become particularly important when the target feedstocks are unconventional, heterogeneous (e.g., seasonal variation) or inhibitory. Classical strain engineering can address individual bottlenecks, but it is often slow and constrained by incomplete knowledge of the genetic determinants. More flexible approaches are needed to diversify and adapt host genomes in parallel with enzyme engineering. Synthetic genomics provides a route to engineer such adaptive capacity directly into microbial chromosomes (James et al., 2025; Schindler, 2020; Schindler et al., 2018). For instance, the Synthetic Yeast Genome Project (Sc2.0) (Erpf et al., 2025; Richardson et al., 2017; Schindler et al., 2024) has implemented an accelerated chassis optimisation tool termed SCRaMbLE (Synthetic Chromosome Rearrangement and Modification by *LoxP*-mediated Evolution) to induce genome-wide structural diversity upon Cre recombinase expression (Box 1; (Dymond et al., 2011)). By producing combinatorial genome rearrangements, SCRaMbLE can rapidly create phenotypic diversity that can be selected under defined environmental or metabolic pressures (Blount et al., 2018; Gowers et al., 2020; Lu et al., 2025). This capability makes synthetic yeast a compelling chassis for exploring how genome-enabled adaptation can be coupled to enzyme discovery and biocatalyst optimisation. In the context of biopolymer valorisation, synthetic-genome-enabled chassis improvement could help identify host configurations that better support enzyme display, substrate tolerance, carbon utilisation or production of value-added compounds. These functions could be growth-coupled (Bushin et al., 2026; Schulz-Mirbach et al., 2026), in which case a large population generated by SCRaMbLE could face competition to select those with increased propagation rates. Surface display is therefore key to stringent selection by tethering enzymes to their host cell; it directly links phenotype to genotype, a link lost with secreted enzymes, whose activity can occur away from the producing cell.

Here, we present an integrated platform for the discovery, evolution, and characterisation of enzymes for microbial biopolymer upcycling, focusing on small laccases from prokaryotes which have remained largely overlooked despite their tolerance to harsh conditions and potential for lignin degradation (Guan et al., 2018; Majumdar et al., 2014). We combine sequence-based discovery, modular yeast surface display, directed evolution, and synthetic yeast genome-enabled chassis improvement to couple enzyme discovery and engineering with host optimisation. We compare natural candidates and an engineered variant within a unified workflow and prove their action on lignin through HPLC-MS analysis. This approach provides a scalable framework for advancing oxidative biocatalysts and establishes laccases as tractable entry points for developing microbial systems for biopolymer valorisation.

### Box 1 SCRaMbLE: Rapid genome evolution via structural variation in synthetic yeast

The Synthetic Yeast Genome Project (Sc2.0) is an international effort to redesign and construct a synthetic genome of *Saccharomyces cerevisiae* (Richardson et al., 2017). Each synthetic chromosome is synthesized in a separate strain with the aim of retaining normal cellular functions but incorporate defined design changes, including removal of transposable elements and selected repetitive sequences, recoding of TAG stop codons, relocation of nuclear transfer RNA genes and insertion of sequence tags that distinguish synthetic from native DNA. A central innovation of Sc2.0 is SCRaMbLE (Synthetic Chromosome Rearrangement and Modification by *LoxP*-mediated Evolution), an inducible system for generating structural variation across synthetic chromosomes (Cheng et al., 2024; Dymond et al., 2011; Shen et al., 2016).

SCRaMbLE relies on symmetrical 34-base-pair Cre recombination sites (*loxPsym*; 5′-ATAACTTCGTATAATGTACATTATACGAAGTTAT-3′) that were placed downstream of nearly all non-essential genes. Expression of Cre recombinase induces recombination between *loxPsym*, resulting in deletions, inversions, duplications and translocations. Multiple events can occur in the same cell, creating complex combinations of changes in gene copy number, order and orientation. In contrast to targeted genome editing, SCRaMbLE does not create a predetermined genotype; it generates large libraries of structurally diverse genomes from a single starting genotype tha t can be screened or selected for useful phenotypes (Dymond et al., 2011; Shen et al., 2016).

Early studies established that Cre induction could remodel synthetic chromosome segments and generate variation in growth and stress-response phenotypes (Ma et al., 2019; Ong et al., 2021). Subsequent work showed that SCRaMbLE could operate across multiple synthetic chromosomes and in haploid, diploid and hybrid backgrounds (Jia et al., 2018; Shen et al., 2018), while reporter systems and long-read sequencing improved the recovery and reconstruction of rearranged genomes (Luo et al., 2018). SCRaMbLE has also been applied to metabolic and industrial strain engineering. Rearrangement of synV (synthetic chromosome V) produced strains with improved violacein and penicillin production and enhanced xylose utilisation (Blount et al., 2018). In a betulinic-acid-producing strain, screening approximately 1,000 SCRaMbLE variants identified isolates with two-to sevenfold higher titres, with long-read sequencing linking improved production to distinct structural rearrangements (Gowers et al., 2020). More recent work has combined iterative SCRaMbLE, fluorescence-activated enrichment and pooled long-read sequencing to optimize synthetic modules and examine how repeated rounds of genome restructuring shape cellular performance (Lu et al., 2025). Importantly, SCRaMbLE is not entirely random: as with any genome-engineering method, its outcomes are shaped by inherent biases, including variation in the chromatin accessibility of individual *loxPsym* (Lindeboom et al., 2024; Zhou et al., 2023).

The main challenge is no longer generating diversity but efficiently screening it and linking phenotype to genotype. Many rearrangements are neutral or deleterious, and beneficial traits can result from interacting changes. Nevertheless, SCRaMbLE provides a powerful route to optimize the host-genome context around engineered pathways.

## Results

### A modular yeast surface-display platform enables enzyme benchmarking and directed evolution

Yeast surface display is a powerful platform for selecting, characterising and engineering enzymes, as it physically links the phenotype of a displayed enzyme to a genotype (Teymennet-Ramirez et al., 2021). In addition, surface-bound enzymes have advantages as they provide catalytic products near the displaying cell, potentially improving utilisation in downstream steps. Establishing display systems with functionally exposed enzymes can be challenging, as multiple variants may affect optimal display. With this in mind, we expanded our previously published modular Golden Gate assembly system (de Vries et al., 2024) by generating a dedicated part library for yeast surface display applications (Figure 1A, Table S1). As a proof of concept, we selected a well-studied small laccase from *Amycolatopsis* sp. (ATCC 39116/75iv2), previously proven to enhance lignin breakdown (Majumdar et al., 2014; Singh et al., 2017; Vuong et al., 2021), hereafter sLac_Amyco_. We displayed sLac_Amyco_ on the yeast cell surface to (i) establish a colorimetric assay for assessing laccase activity with the aim of using this system for directed evolution and (ii) characterise our Golden Gate assembly parts. This will enable the selection of improved variants after rounds of directed evolution, while providing a platform for the experimental characterisation of computationally mined laccases.

**Figure 1.**
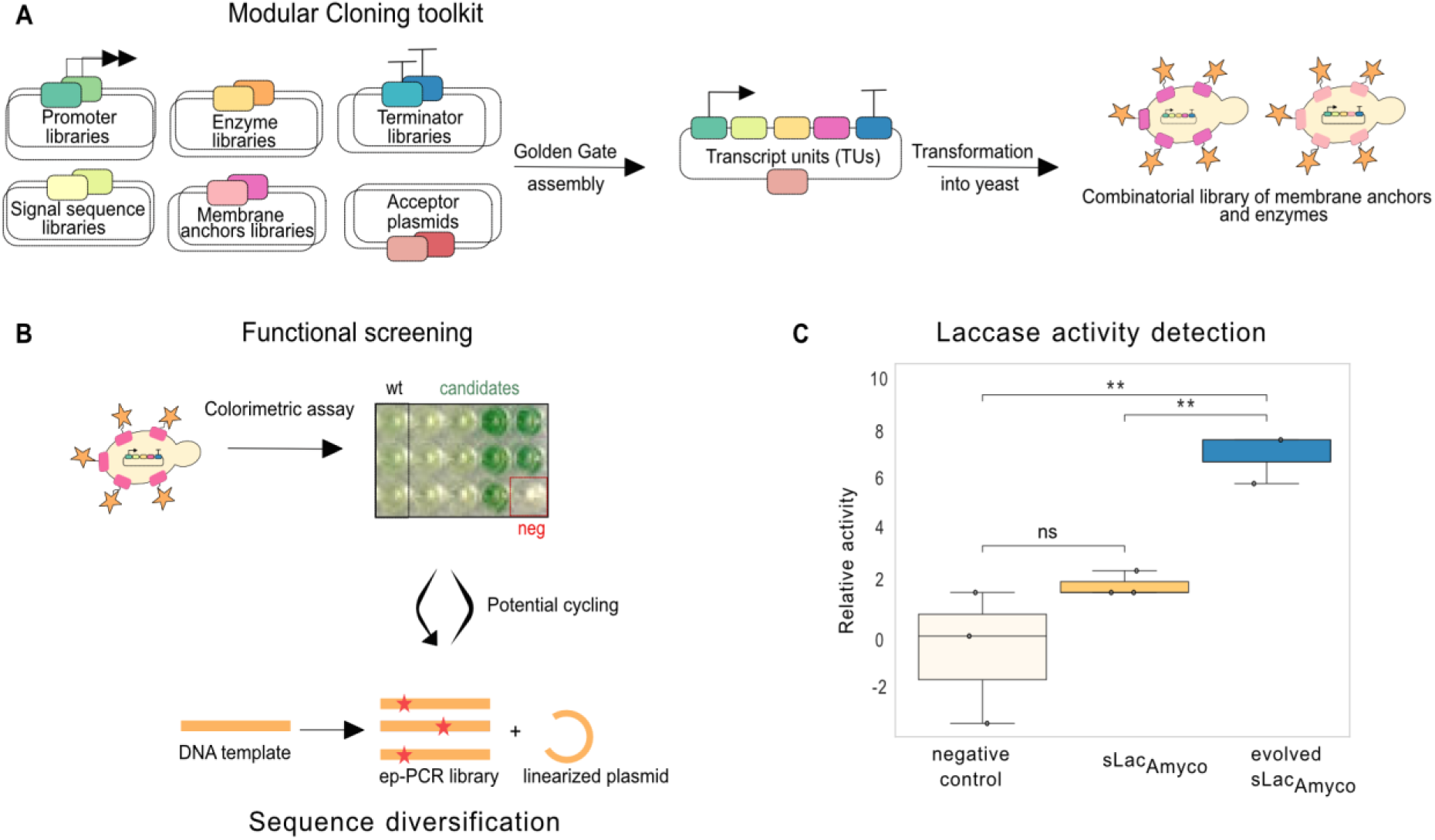
Yeast surface display toolkit for enzyme activity detection and directed evolution. **(A)** Modular Cloning toolkit for yeast surface display. **(B)** Screening workflow combining the colorimetric ABTS assay with sequence diversification via error-prone PCR (ep-PCR) for random mutagenesis. **(C)** Boxplot showing the relative activity of evolved sLacAmyco to sLacAmyco. Whiskers indicate min and max, points show individual replicates (n = 3). One-way ANOVA, F = 17.18, P = 0.0033; Kruskal-Wallis, P = 0.0349. Pairwise comparisons: two-sided Student’s t-test with Holm–Bonferroni correction. Evolved sLacAmyco vs. sLacAmyco, P = 0.0013; evolved sLacAmyco vs. negative control, P = 0.0088; negative control vs. positive control, P = 0.17. *P < 0.05, **P < 0.01, ***P < 0.001.

Enzymatic activity of laccases can be quantified using the colorimetric ABTS (2,2′-azino-bis(3-ethylbenzothiazoline-6-sulfonic acid)) assay (Johannes & Majcherczyk, 2000). The oxidation of ABTS by laccase generates the visibly coloured ABTS cation radical (Figure 1B), which can be measured spectrophotometrically by monitoring its absorbance at 418 nm (Pardo et al., 2013). We purposely selected a colorimetric assay with the vision to enable high-throughput screening of engineered and mined laccases. With this vision in mind, we explored whether laccase activity could be monitored directly on agar plates using a colorimetric readout in combination with high-density spotting of yeast strains. However, despite testing agar-plate-based conditions and different reported colorimetric assays such as syringaldehyde and sinapic acid (Pardo et al., 2013), we were unable to establish a robust assay for reliable screening and selection. Therefore, subsequent activity assessments are based on liquid colorimetric ABTS assay measurements. Notably, the ABTS assay only indicates enzymatic ABTS oxidation and does not indicate polymer (e.g., lignin) degradation, which we later investigated via HPLC-MS analysis of laccase-catalysed kraft-lignin depolymerisation.

To establish our experimental system, we displayed sLac_Amyco_ with different membrane anchor proteins: AGA2, CIS3, CWP2 and TIP1 under control of the inducible gal10 promoter (genetic parts are listed in Table S1 and sequences are provided in the Supporting Data). For yeast surface display we used BY4741 as a wild-type reference strain in which SED1 was deleted, resulting in strain SLy0475. SED1 encodes a highly abundant cell wall glycoprotein and its deletion was previously shown to be beneficial for yeast surface display (Kotaka et al., 2010). Our combinatorial modular approach showed that activity varied considerably across the four anchor-fused sLac_Amyco_: N-terminal fusion of sLac_Amyco_ to the CIS3 anchor (SLy0519) gave the best performance in the ABTS assay, whereas the remaining three anchor proteins showed no measurable activity in the ABTS assay (data not shown). Subsequently, we used our established system with sLac_Amyco_ fused to CIS3 to implement a round of directed evolution using established error-prone PCR (ep-PCR) protocols (Mate et al., 2010). After screening ∼600 candidates in the first generation of directed evolution, we retrieved a variant with 1.65-fold higher activity than the parental sLac_Amyco_ (Figure 1C). The resulting variant contains the mutation D230Y. Two-domain, or ’small’, laccases such as sLac_Amyco_ differ from the more widely studied three-domain laccases, as they contain only two cupredoxin domains and form either homodimers or homotrimers. Strikingly, the D230Y mutation is located at the monomer interface (Figure S1), potentially stabilising subunit interactions. This may account for the increased consistency in the measurements of the evolved sLac_Amyco_ variant relative to its parental version.

### Sequence mining identifies divergent unexplored laccases for functional screening

Having established the yeast surface display platform, our aim was to expand the range of enzymes available for lignin depolymerisation. To this end, we mined publicly available sequence databases for remote homologues of small bacterial laccases with catalytic potential. Laccases [EC:1.10.3.2] are divided into two distinct orthologous groups in KEGG (Kanehisa & Goto, 2000) reflecting their evolutionary and biological differences. Canonical eukaryotic laccases, including the well-characterized plant representatives (Cai et al., 2006) are assigned to orthogroup K05909 (hereafter referred to as *eukaryotic laccases*). In contrast, prokaryotic laccases, including archaeal representatives (Uthandi et al., 2010) such as LccA from *Haloferax volcanii,* belong to orthogroup K00421 (hereafter referred to as *prokaryotic laccases*). To broaden our knowledge of characterized small laccases beyond enzymes described to date, we conducted a deep homology search to identify potential remote homologues (Figure 2A). Building on KEGG laccase classification and using the experimentally characterized lignin-degrading sLac_Amyco_ as query, we screened a curated dataset of 19,642 reference genomes and 149,842 metagenomes (Rodriguez Del Rio et al., 2024) spanning the search over 40 million gene predictions for undescribed remote homologues of *prokaryotic laccases*. We identified three sets of enzyme candidates. The first set contains 95 sequences that share at least 25% sequence similarity with *prokaryotic laccases*; this set is designated sLac01. The second set, sLac02, contained 182 sequences that, unlike sLac01, showed sequence similarity of at least 25% to proteins annotated as hydroxylamine dehydrogenases [EC:1.7.2.6] (Hommes et al., 2001) within orthogroup K10535. Additionally, we identified 277 putative laccases, sLac03, that share similarity with *eukaryotic laccases*. However, this set of sequences was excluded from further analysis as our focus was on identifying remote homologues of *prokaryotic laccases*.

**Figure 2.**
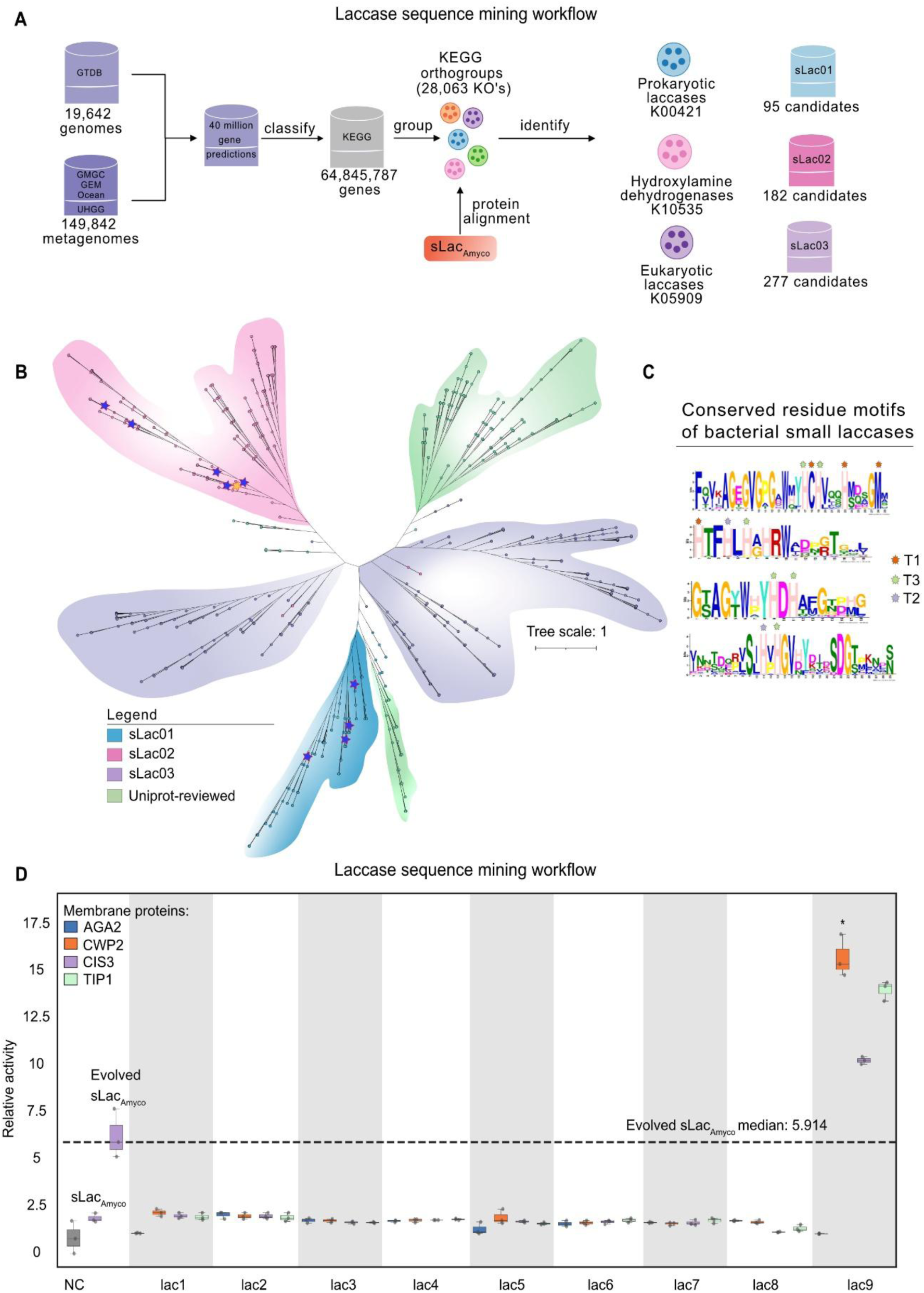
Mining public databases for unexplored laccases and activity assessment of selected candidates. **(A)** Data mining workflow of genome and metagenome curated repositories. After coding sequence identification and classification according to KEGG orthogroups, a deep homology search using sLacAmyco as query generated three distinct sets of sequences with similarity to *prokaryotic laccases* (sLac01, in blue), *hydroxylamine dehydrogenase*s (sLac02, in pink) and *eukaryotic laccases* (sLac03, in purple). **(B)** Phylogeny of the three sets of putative laccases with Uniprot-derived curated multicopper oxidoreductases. After sequence, structure and metabolic potential assessment, nine laccases (marked with blue stars) were selected for experimental characterization, slacAmyco is highlighted by a yellow star. **(C)** MEME motifs of the conserved residues forming the copper cluster in laccases. Stars in distinct color indicate the respective residues forming either of the T1, T2 and T3 copper centers. **(D)** Relative activity of small laccases (slac), each fused to one of four anchor proteins (AGA2, CWP2, CIS3, TIP1), compared with evolved sLacAmyco. Box plots show median and interquartile range; points show individual replicates (n = 3). One-way ANOVA, F = 260.133, P < 0.0001; Kruskal-Wallis, P < 0.0001. Levene’s test indicated unequal variances across groups (stat = 1.733, P = 0.0207); pairwise comparisons therefore used two-sided Welch’s t-test with Holm–Bonferroni correction across 38 comparisons, each candidate versus the evolved sLacAmyco. *P < 0.05, **P < 0.01, ***P < 0.001. Eight out of nine tested candidate laccases showed no activity while sLacSspir (lac9) showed a drastic increase in the ABTS assay compared to the evolved sLacAmyco. lac1: *H. sedimenticola*; lac2: *A. bryophytorum*; lac3: SIRX01; lac4: *M. violae*; lac5: *Methylomicrobium sp016745425*; lac6: *P. resinovorans*; lac7: *A. xinjiangensis*; lac8: *P. flavigriseum*; lac9: *S. spiralis*.

To further assess the extent to which small laccases are distributed across taxonomic lineages and to pinpoint their potential functional orthologues, we queried sLacAmyco against the reviewed UniProtKB (Swiss-Prot) database (release 2025_04) (UniProt, 2007). We obtained 192 curated sequences that fall into the multicopper oxidase protein family, ID: IPR045087. The curated sequences mainly belong to eukaryote-derived multicopper oxidases, oxidoreductases and laccases from the phyla Streptophyta, Ascomycota and Basidiomycota. In addition, from this set, 45 prokaryotic sequences, belonging to Pseudomonadota with various functions, from laccase to copper-containing nitrite reductases and cell-division-related proteins were obtained. To understand the evolutionary relationships between the UniprotKB sequences and identified sequence groups (sLac01-sLac03) a phylogenetic tree was constructed (Figure 2B). Most of the sequences derived from sLac03 group with eukaryotic multicopper oxidases. The sequences in this group mainly belong to the Proteobacteria (68%), which show a preference for three cupredoxin domains (Figure S2).

In this study, we focus on two-domain laccases that contain both cupredoxin domains (Cu-oxidase-2 and Cu-oxidase-3). The sLac01 sequences predominantly belong to the phyla Actinomycetota and Halobacteriota (Figure S2). The fact that these distant homologues cluster with bilirubin oxidases and O-aminophenol oxidases, which are involved in pigmentation, suggests that the sLac01 sequences may have a biological function distinct from that of “canonical” small laccases such as sLac_Amyco_. The sLac02 sequences belong mostly to the phyla of Actinomycetota and Proteobacteria (Figure S2). From an ecological perspective, Actinomycetales are ubiquitous in soil ecosystems, playing a general role in litter decomposition and carbon cycling. They are able to secrete a variety of enzymes, including lignocellulolytic enzymes such as xylanases and cellulases, as well as laccases (Lewin et al., 2016). This reinforces the potential of these remote homologues as candidate small laccases with lignin-degrading activity.

sLac01 and sLac02 sequences were manually curated via motif detection and Pfam domain identification, retaining those with conserved laccase active-site residue motifs (Figure 2C). This resulted in 34 putative sLac01 and 81 sLac02 sequences. Structure conservation analysis, Pfam domain identification, and AlphaFold structural predictions of candidate sequences containing active-site motifs were compared to the sLac_Amyco_ (PDB: 3TA4) and reference laccase structures (Figure S3 and S4). The metabolic potential of source organisms was assessed by identifying protein families involved in both lignocellulose degradation and lignin-monomer assimilation using Pfam domain identification (Figure S5 and S6). Together, this information guided the selection of nine candidate small laccases (Table S2) for gene synthesis and functional testing using our modular yeast surface display and the ABTS-based colorimetric assay.

### Surface-display characterisation reveals functional variation among laccases

The nine selected laccase sequences were computationally codon-harmonised *S. cerevisiae* and selected type IIS recognition sites were removed (Claassens et al., 2017; Zulkower & Rosser, 2020). Using our modular Golden Gate toolkit all nine selected laccases were fused N-terminally to the four different anchor proteins, resulting in a total of 36 expression constructs (pSL1142 to pSL1176). The plasmids were transformed into the BY4741 expression platform strain (SLy0475). In the first screening, their ability to oxidise substrates was tested using the ABTS assay, using the evolved sLac_Amyco_ (pSL1179) as reference. Eight of the nine sequences did not show activity in the ABTS assay (Figure 2D). Yet, one laccase identified in our mining approach, from *Streptomyces spiralis* and fused to CIS3, showed 5.2-fold higher activity than the already evolved sLac_Amyco_. This laccase is hereafter termed sLac_Sspir_. The enzyme showed even 7.7- and 6.8-fold higher activities when fused with CWP2 and TIP1 anchor proteins, respectively compared to the evolved sLac_Amyco_. These results show that the way a protein is displayed can significantly affect its activity. This emphasises the importance of adopting a modular approach, which enables the rapid testing of various protein display combinations.

Subsequently, we performed a time series experiment to evaluate the ability of the evolved sLac_Amyco_ and the mined sLac_Sspir_ (fused to CIS3 for comparison purposes) to depolymerise Kraft-lignin. Yeast cells expressing and surface displaying sLac_Amyco_ and sLac_Sspir_, as well as the control strain (SLy0546), were incubated with Kraft-lignin and ABTS, which acts as a redox mediator to promote lignin degradation (Figure 3A). Samples were collected at the starting time point (0 h) and at six subsequent time points (2 to 72 h) and analysed using HPLC-MS. Our results show that the depolymerisation profiles of sLac_Amyco_ and sLac_Sspir_ differ with regard to the products generated and their temporal effects on lignin degradation (Figure 3B, S7, and Table S3), which potentially implies that the tested laccases have different modes of action.

**Figure 3.**
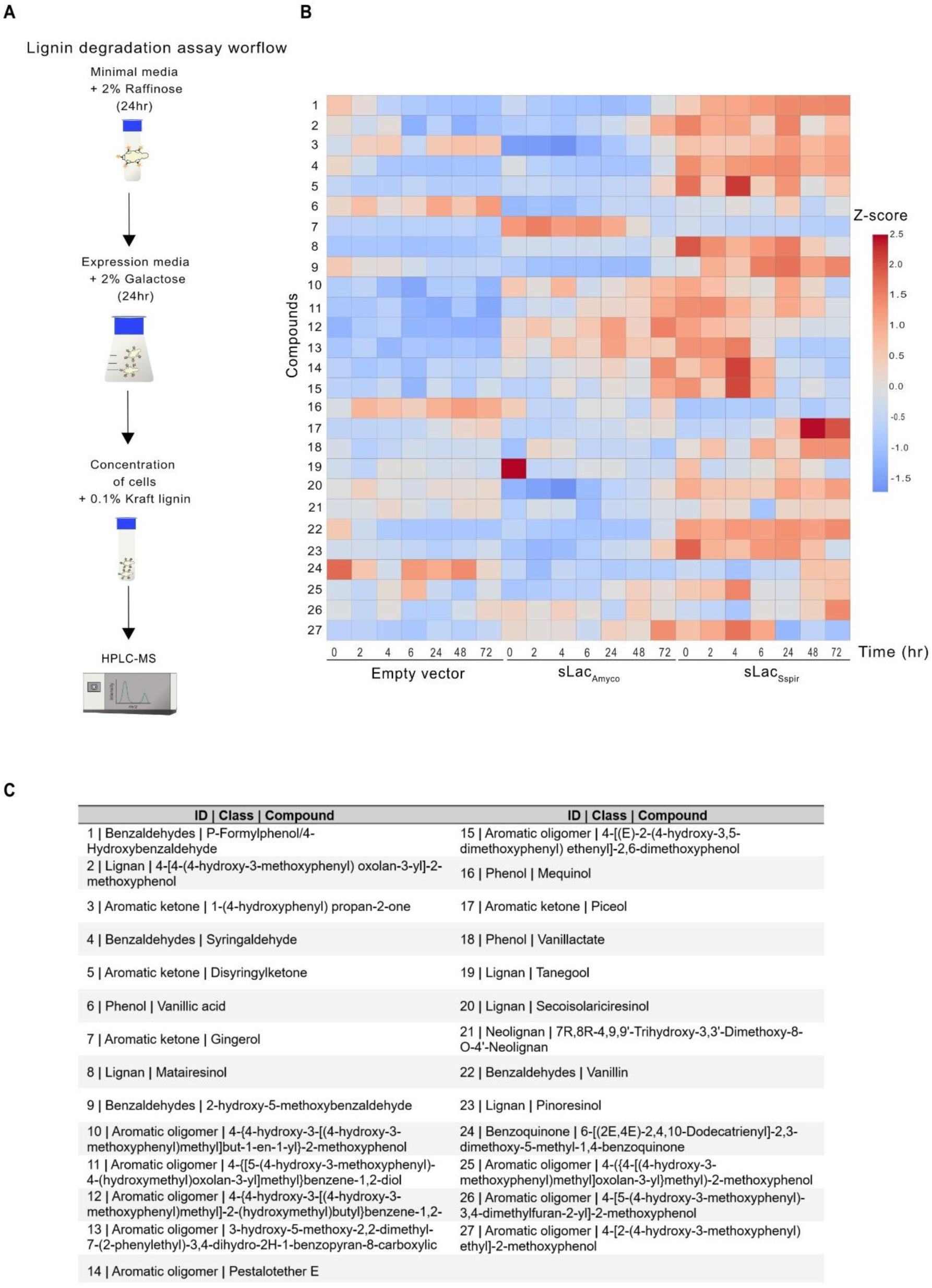
HPLC-MS analysis provides insights into lignin depolymerisation profile of laccases. **(A)** Yeast strains were grown in minimal medium to establish the culture, then transferred to expression medium for laccase surface display. Cells were subsequently harvested, concentrated and subsequently incubated with 0.1% (w/v) kraft-lignin for a depolymerisation analysis. Samples were collected over a time course (0 to 72 h) and analysed by HPLC-MS to identify and quantify the resulting depolymerisation compounds. **(B)** Heatmap of z-score-transformed signal intensities of lignin depolymerisation products (numbers on left) generated by sLacAmyco and sLacSspir over time. Signal intensities for each detected compound were z-score normalised to enable comparison across compounds and time points; colour scale represents relative abundance (z-score). n = 3 biological replicates. Box plots for each compound across samples and time points, along with the statistical analysis are shown in Figure S8. Figure S9 provides an overview of the effects of treatment, incubation time and compound signal intensity. **(C)** Table listing the detected compounds of (B).

The results of sLac_Sspir_ indicate, in contrast to sLac_Amyco_, a higher abundance of monomeric aromatic compounds of the benzaldehyde class, such as vanillin and 4-hydroxybenzaldehyde. These compounds are widely observed after laccase-mediated lignin depolymerisation (de Jong et al., 2020; Sainsbury et al., 2013; Vignali et al., 2022; Yang et al., 2019). Notably, the presence of natural redox mediators such as syringyl-derived compounds affects the lignin transformation process (Chen et al., 2021). The presence of syringaldehyde (compound 4 in Figure 3) in sLac_Sspir_-treated samples may contribute to the acceleration of the laccase-catalysed reaction, enhancing lignin depolymerisation. Interestingly, a depletion of (4) was previously reported for sLac_Amyco_ (Singh et al., 2017), which could suggest a rapid oxidation of (4) by sLac_Amyco_ and therefore reflects the different modes of action among the tested laccases. Additionally, lignan synthesis has been reported to be facilitated by small laccases (e.g., in *Streptomyces coelicolor* (Nemadziva et al., 2018)). The abundance of lignan-related compounds found in the lignin depolymerisation process by sLac_Sspir_ could indicate the dimerisation of eugenol or coniferyl alcohol units into pinoresinol and associated compounds (Figure 3: 2, 8, 14, 13, 12, 11, 10 and 27). sLac_Amyco_ generates lignan-related compounds too (Figure 3: 10, 11, 12, 13, 14), but with lower abundance.

Our results support the concept that lignin degradation can be achieved using yeast-surface-bound bacterial laccases. Previously, this process had only been demonstrated for fungal laccases (Bertrand et al., 2016; Fernández-Sandoval et al., 2026). In a recent study, the directed evolution of the AGA2 anchor was shown to improve the activity of a fungal laccase (Teymennet-Ramírez et al., 2026).

Having demonstrated the effectiveness of sequence-based discovery in conjunction with yeast surface display for experimental validation and directed evolution for improving enzymes, we aimed to investigate how the choice of expression host influences laccase activity, particularly in terms of its impact on protein secretion, processing, and exposure at the yeast cell wall. As the synthetic yeast genome project has now made a full panel of synthetic chromosomes available (Archer et al., 2026; Richardson et al., 2017), we transformed our sLac_Sspir_ expression plasmid into various strains and used the ABTS assay to quantify improvements based on strain background. This approach also enabled us to exploit SCRaMbLE, a system that induces rapid, controlled genome rearrangements in these strains, to further explore the effects of the genome on enzyme performance (Dymond et al., 2011; Shen et al., 2016).

### Synthetic yeast strain backgrounds shape laccase display and activity

In order to improve the background of the yeast expression strain, we decided to carry out the transformation of the plasmid pSL1160 encoding sLac_Sspir_ for surface display in all published single synthetic yeast chromosome strains (Annaluru et al., 2014; Blount et al., 2023; Foo et al., 2023; Goold et al., 2025; Lauer et al., 2023; Luo et al., 2023; McCulloch et al., 2023; Mitchell et al., 2017; Richardson et al., 2017; Shen et al., 2023; Shen et al., 2017; Williams et al., 2023; Wu et al., 2017; Xie et al., 2017; Zhang et al., 2023; Zhang et al., 2017; Zhou et al., 2024). The only exceptions, with two synthetic chromosomes, are the synI/III and synI-III strains which have either the separate versions of both synthetic chromosomes (as in synI/III), or a fused version (synI-III) to prevent the loss of the smallest chromosome and to obtain a 16-chromosome karyotype in the final Sc2.0 strain which will contain the 17^th^ chromosome, the tRNA neochromosome (Luo et al., 2023; Richardson et al., 2017; Schindler et al., 2023).

After transformation of pSL1160 into the different strain backgrounds we performed the ABTS assay to determine the enzyme performance in comparison to the BY4741 reference expression platform strain containing pSL1160 (SLy0766) (Figure 4A). For most of the synthetic strains, except synX and synXVI, we observe no effects on cell growth when expressing the construct with sLac_Sspir_ (data not shown). Interestingly, we observed strong differences between the strains even without any additional genome modifications, which highlights the importance of the strain background (Figure 4A). In particular, synVII has a ∼7.8-fold higher performance in the ABTS assay compared to BY4741 (Sly0766). Based on our observation we looked into the published transcriptome and proteome data of synVII (Shen et al., 2023). At the transcriptome level, three transmembrane proteins involved in cell wall stability, organisation, and remodelling (YGR023W, YKL163W, YLR194C) are upregulated, along with a small heat shock protein with chaperone activity (YDR171W), while a transmembrane protein of the major facilitator superfamily (YNL065W) is downregulated. At the proteome level, reduced abundance of two transmembrane proteins (YGL053W, YML123C) and one cell wall mannoprotein (YKL096W) is observed. The differential expression and protein abundance may affect cell wall remodelling and plasma membrane dynamics, resulting in the observed difference in laccase activity in synVII in the ABTS assay. While we do not anticipate any difference in catalytic function, we speculate that there may be a difference in the amount or clustering of the displayed enzyme.

**Figure 4.**
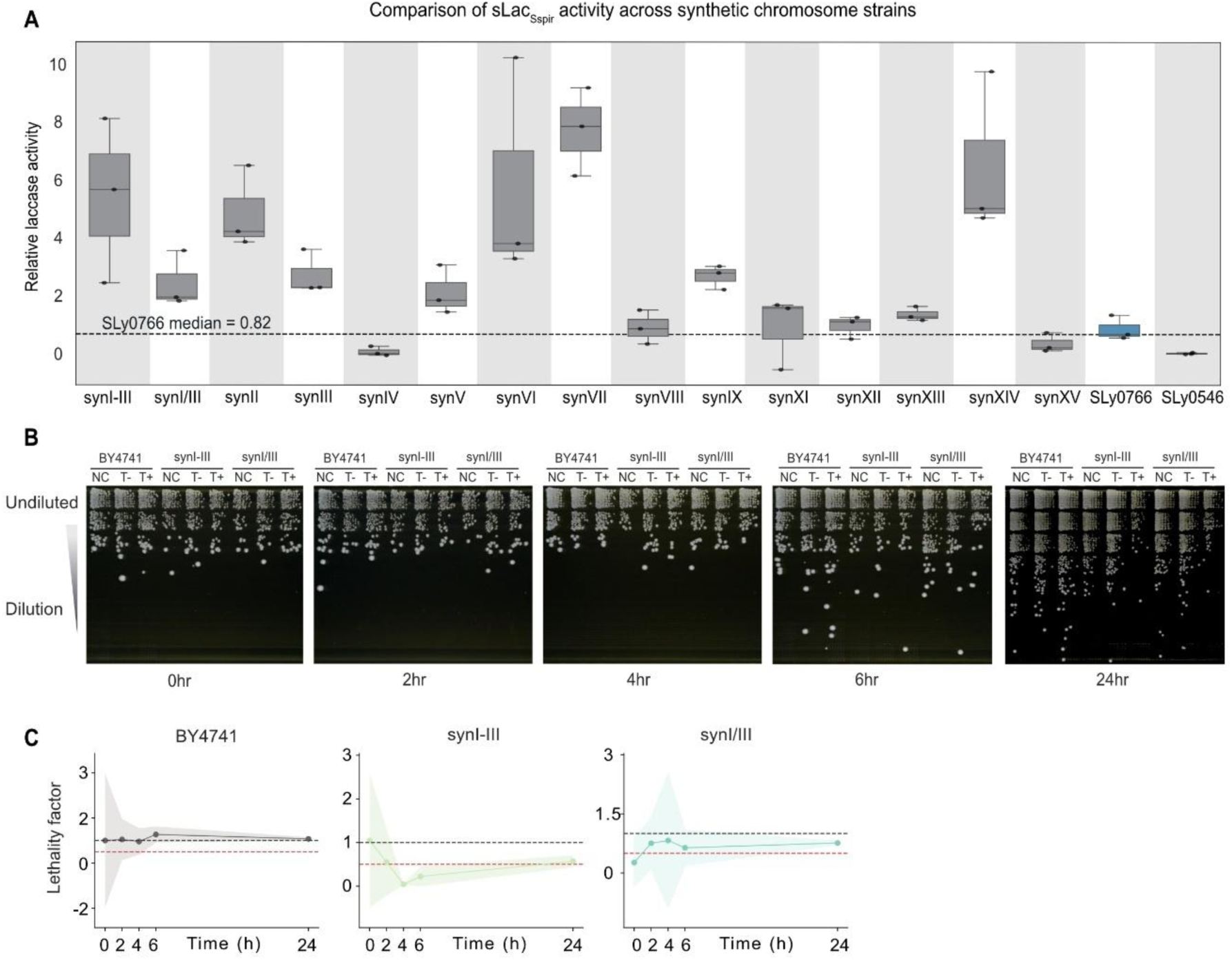
Relative activity across synthetic chromosome yeast strains of sLacSspir and determination of the optimal SCRaMbLE time point. **(A)** Boxplot showing relative activity of sLacSspir, measured by the ABTS assay, in the indicated synthetic yeast strains, normalised to the SLy0776 control strain (n = 3). Statistical significance was assessed by the Kruskal-Wallis test followed by Dunn’s multiple comparisons test against SLy0776; no significant differences were detected between any synthetic strain and the control (p > 0.05 for all comparisons). synX and synXVI were excluded from this assay based on the observed growth phenotype. **(B)** Representative, exemplary Cre-induced killing assay. 10-fold dilution series of wild-type (BY4741), synI-III and synI/III with empty plasmid (NC), and uninduced (T-) and induced (T+) Cre expression plasmid (pSL0217 or pSCW11-CreEBD). **(C)** Ratio of normalised colony forming units to determine the lethality factor in the Cre-induced killing assay for synI-III and synI/III in comparison to the wild-type. Black dotted line indicates 1, red line 0.5, shaded area in each plot represents the standard deviation within the killing assay (n = 3).

synI-III, synVI, and synXIV show an increased activity in the ABTS assay relative to SLy0766 with 5.5, 5.9, and 6.6-fold improvement, respectively. However, the measurements show variability among the biological replicates (Figure 4A). synV, synIX, synXII, and synXIII showed a 2.26, 2.82, 1, and 1.5-fold increased activity compared to SLy0766, respectively with lower variability between the individual measurements. Overall, our results show that the underlying genotype has an impact on monitored activity in yeast surface display. The strains used have the same BY4741/BY4742 background and differ only by one synthetic chromosome (with the exception of synI/III and synI-III). Thus, the overall gene content is highly similar and SED1 is only deleted in the control strain SLy0766.

The observation of different activity profiles of sLac_Sspir_ in the individual synthetic chromosome strains led us to ask if SCRaMbLE would further improve laccase activity, as well as identify genes involved in cell wall composition, protein trafficking and folding that have an impact on the surface-displayed sLac_Sspir_.

### SCRaMbLE enables rapid chassis improvement for enhanced enzyme performance

Sc2.0 SCRaMbLE-based genome evolution is an outstanding tool for genome diversification; it rapidly creates a complex population from a single genotype (Dymond et al., 2011; Shen et al., 2016). However, due to its random recombination events, it can also result in the loss of essential genes leading to cell death. We exploited this feature to determine the optimal induction time for Cre recombinase for each synthetic chromosome strain (Figure 4B-C and Figure S10-S11). We defined the optimal time point for SCRaMbLE at which around 50% cell death after induction of Cre occurs. Given this, we can assume that the majority of colonies obtained should have Cre-mediated recombination events. Therefore, prior to SCRaMbLE, we performed a systematic assay to identify this time point with cell lethality defined as a reduction in colony-forming units (CFUs). Despite our lethality-assay having a broad standard deviation, we did not observe any correlation with regard to chromosome length or the number of essential genes encoded. Based on these results (Figure 4B-C and Figure S10-S11), we selected 6 h as the induction time used in this study for all synthetic chromosome strains. We concluded that, in the absence of a stringent selection scheme, it is important for any SCRaMbLE experiment to perform a CFU comparison of the induced and uninduced culture.

Having determined the optimal induction time for SCRaMbLE, we conducted genome diversification on all synthetic chromosome strains (except synX and synXVI, data not shown) containing sLac_Sspir_ (pSL1160) and expressing Cre recombinase (pSL0217 or pSCW11-CreEBD). We randomly selected 48 candidates for each synthetic chromosome and performed the ABTS assay. We were interested in strains showing improved ABTS oxidation in order to analyse their genomes for beneficial alterations for sLac_Sspir_ display (Figure 5A). Additionally, we assumed that candidates showing little or no ABTS oxidation could provide valuable information regarding genotype alterations that reduce or diminish sLac_Sspir_ display. This could validate our mechanistic understanding of protein trafficking and display and could provide hints for genes whose overexpression may contribute to better performance in yeast surface display. Of the 15 synthetic strains subjected to SCRaMbLE we found candidates with relevant changes in sLac_Sspir_ activity across synI/III, synVI, synVII, synVIII and synIX. We selected 13 strains and performed long-read Nanopore sequencing to connect phenotype and genotype. No aneuploidies were detected based on sequencing data (Figure S12). However, whole-genome duplication, as previously observed, cannot be ruled out (Lindeboom et al., 2024).

**Figure 5.**
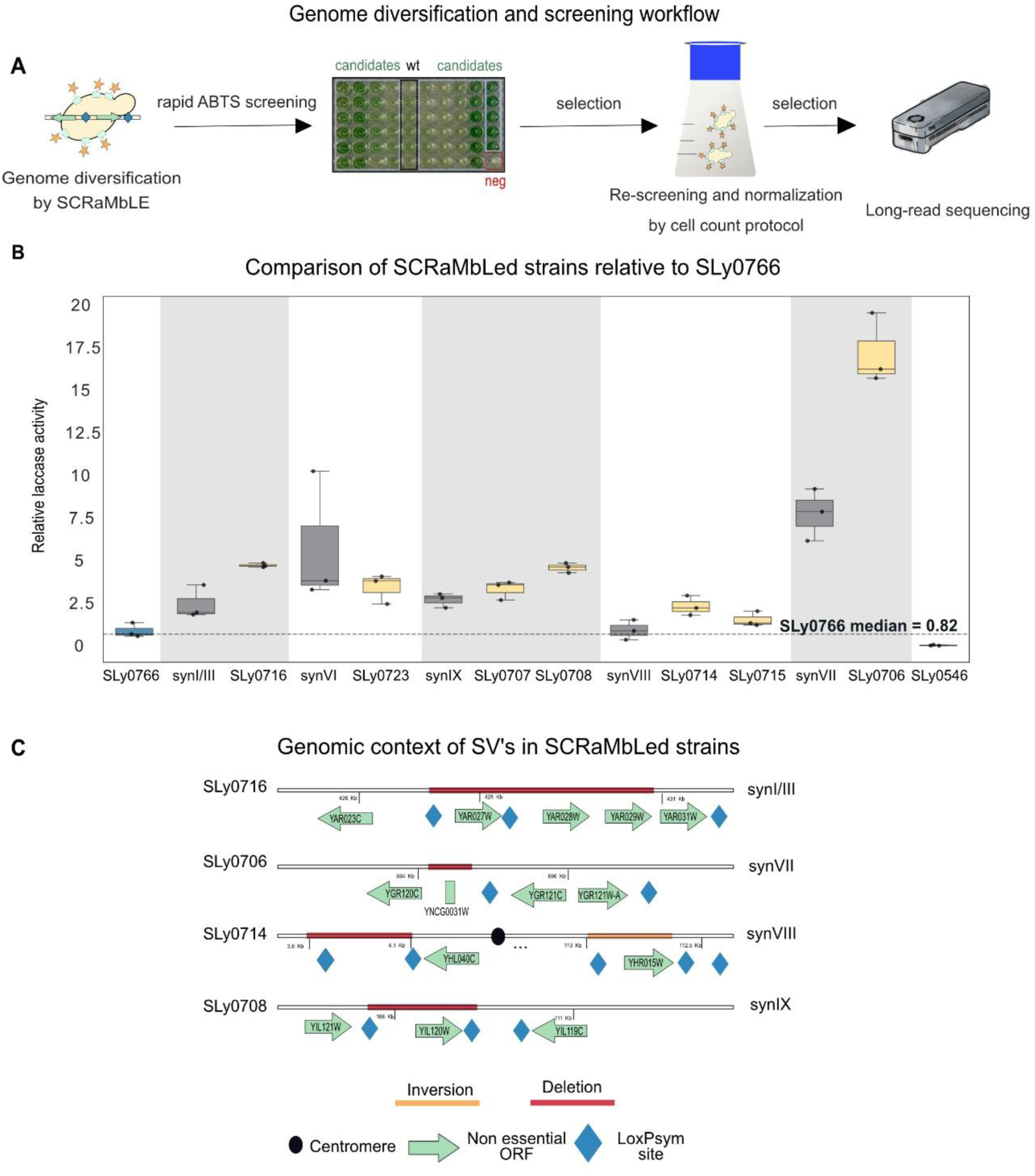
SCRaMbLE generates a diverse population with candidates showing improved yeast surface display properties. **(A)** Workflow for genome evolution using SCRaMbLE in synthetic yeast strains. **(B)** Boxplot of relative activity of selected SCRaMbLEd strains in comparison to the parental strain and the SLy0766 reference strain (set to 1); n = 3. Statistical significance was assessed by the Kruskal-Wallis test followed by Dunn’s multiple comparisons test against SLy0766; no significant differences were detected between any synthetic strain and the control (p > 0.05 for all comparisons)**. (C)** Detected SVs which are most likely caused by Cre recombinase in selected SCRaMbLEd strains. Plots for whole genome coverage and dot plots for the rearranged chromosomes are provided in Supporting Figures S12-S13.

Overall, the SCRaMbLEd strains showed improved activity when compared to the reference strain SLy0766 (Figure 5B). The SCRaMbLEd synVII candidate, SLy0706, showed the most significant increase in ABTS activity, at 17.3-fold. This was followed by candidate SLy0716 (synI/III) with a 4.9-fold increase and the synIX candidates SLy0707 and SLy0708, which showed a 3.4- and 4.7-fold increase in the ABTS assay, respectively. We used Nanopore sequencing to link the observed phenotype to its genotype. As hypothesised, we found genotype alterations that cause changes in the regulation or expression of genes involved in cell wall composition, trafficking and folding of proteins caused by structural variations (SV). Those alterations most likely do not change the activity of the displayed laccase but may increase the amount of displayed protein on the cell surface due to multiple causes (e.g., more efficient trafficking).

We observed the deletion of membrane-associated proteins when we analysed the long-read sequencing data. In SLy0716 (synI/III) the genes YAR027W and YAR028W were deleted, both are transmembrane proteins, members of the DUP240 family. In SLy0708 (synIX) YIL120W, a gene that encodes a multidrug transporter, was deleted. Moreover, SLy0706 (synVII) exhibits a deletion spanning a tRNA gene. While tRNA genes were removed from all synthetic chromosomes, this one was reintroduced into synVII due to observed fitness defects during its characterisation (Shen et al., 2023).

The established workflow now enables a large number of SCRaMbLE experiments to be performed in combination with characterisation of laccase functionality on the yeast surface. By establishing links between phenotypes and genotypes, our understanding of protein trafficking and enzyme display on the yeast surface can be expanded. This should validate existing knowledge but may also reveal new important factors.

## Discussion

### Core achievement: Coupling enzyme engineering with synthetic yeast chassis evolution

This study is the first to combine enzyme discovery, evolution, and synthetic yeast chassis adaptation for functional screening and lignin depolymerisation assessment. We extended our modular cloning toolkit (de Vries et al., 2024) for yeast surface display to enable the functional evaluation of plant biomass-depolymerising enzymes, including previously uncharacterised enzymes obtained from public repositories. We demonstrated that this display system is compatible with directed evolution strategies for improving catalytic performance. Furthermore, by leveraging genome evolution enabled by the Sc2.0 project, we demonstrated that chassis engineering is a promising approach for further increasing catalytic activity. In the future, we intend to use our system to advance biomanufacturing and the circular bioeconomy by increasing the use of sustainable feedstocks and upcycling biological waste streams, such as lignin, by incorporating additional valorisation pathways.

### Enzyme discovery and optimisation

Effective lignin depolymerisation depends on several factors, including the catalyst and the availability of redox mediators. sLac_Amyco_, a bacterial two-domain laccase, has been reported to efficiently depolymerise lignin (Vuong et al., 2021), which is why we used it as a starting point for our study. By mining public databases, we identified a laccase that we termed sLac_Sspir_, which outperformed our evolved sLac_Amyco_ in our yeast surface display. These results suggest that additional high-performing enzymes are likely to remain unexplored in sequence databases, given that we made this observation after screening just nine sequence candidates. With modular synthetic biology tools and the established colorimetric assay on hand, we can now test a large number of sequences and thus expand the enzymatic toolkit for lignin valorisation. We also demonstrate that the discovered laccase and the evolved laccase are capable of depolymerizing kraft-lignin. Interestingly, we observed distinct product fingerprints which likely reflect differences in their catalytic mechanisms and how surface immobilisation affects protein display. As shown for sLac_Sspir_, the choice of the anchor protein directly affects how the enzyme is exposed on the cell surface. Accordingly, the optimal combination of promoter, anchor protein, signal sequence and N-or C-terminal fusion is protein-dependent. Our modular cloning system provides a platform to readily mix and match these elements, enabling combinatorial optimisation of enzyme catalytic activity. Nevertheless, it is important to remember that directed evolution produces what is selected for (Schmidt-Dannert & Arnold, 1999).

When selecting for increased functionality in a surface-display system, the observed mutation may improve the display rather than the catalytic function. This may be the case for sLac_Amyco_ as variants fused to different anchor proteins showed only marginal activity, while the D230Y mutation likely increased monomer stability, favouring a stable homotrimeric configuration. This oligomerisation, rather than a genuine gain in catalytic efficiency, could explain the consistent ABTS oxidation we detected in subsequent activity measurements.

The sequence-based discovery of sLac_Sspir_ once again highlights the abundance of valuable, unexplored enzymes. The described sLac_Sspir_ exhibits closer sequence similarity to hydroxylamine dehydrogenases than to prokaryotic laccases, which may explain why it had not been identified earlier the potential of our method to identify divergent homologues with catalytic activity. Furthermore, this finding could suggest a link with nitrogen metabolism. It has been suggested that small laccases could represent an evolutionary link between three-domain laccases and nitrite reductases (Nakamura et al., 2003). Furthermore, it has been demonstrated that hydroxylamine oxidoreductase from Epsilonproteobacteria acts as a missing link in the evolution of nitrite reductases (Haase et al., 2017). Notably, small laccases and nitrite reductases are able to form homotrimeric structures, which raises the possibility that small laccases could represent a link in the evolutionary trajectory between three-domain laccases and nitrite reductases. We hypothesise that sLac_Sspir_ may possess additional, as yet uncharacterised, activities beyond the laccase-like activity investigated here.

### State-of-the-art chassis engineering using synthetic genomics resources

More broadly, this study provides a route for expanding surface-display-based engineering beyond laccases towards diverse enzyme classes and for combining enzyme evolution with rapid chassis diversification through SCRaMbLE. By linking biocatalyst optimisation to genome-enabled host adaptation, such strategies could support the development of yeast platforms able to utilise unconventional carbon sources and heterogeneous waste streams. Based on their initial characterisation, the synthetic strains individually exhibit wild-type-like fitness under the tested conditions unless stated otherwise (Annaluru et al., 2014; Blount et al., 2023; Foo et al., 2023; Goold et al., 2025; Lauer et al., 2023; Luo et al., 2023; McCulloch et al., 2023; Mitchell et al., 2017; Richardson et al., 2017; Shen et al., 2023; Shen et al., 2017; Williams et al., 2023; Wu et al., 2017; Xie et al., 2017; Zhang et al., 2023; Zhang et al., 2017; Zhou et al., 2024). However, this does not account for all potential molecular phenotypes. These may result in subtle differences that do not necessarily manifest as a monitored phenotype under the tested growth conditions, but which may do so under different conditions or applications of the strains. In our study, we observed that the display of our mined laccase differed between the various tested strains, indicating the importance of the strain background. We did not anticipate that the differences would be so drastic without SCRaMbLEing in the different synthetic chromosome strains. This is an interesting and relevant finding for our current and future studies. Subsequent SCRaMbLE experiments further demonstrate that the genotype can be improved, opening up the possibility of a systematic analysis of which structural variants benefit the surface display. This opens up new possibilities for improving yeast surface display for applications and for gaining a deeper understanding of the underlying molecular mechanisms involved.

### Future directions: Growth-coupled selection

In the future, we will replace indirect colorimetric screening with growth-coupled selection. This will make enzyme performance an explicit determinant of cellular fitness (Bushin et al., 2026; Schulz-Mirbach et al., 2026). Previous studies have demonstrated the effectiveness of growth-coupled approaches in optimising metabolic and enzymatic functions (Blount et al., 2018; Orsi et al., 2021), and this method would facilitate higher throughput, reduce the number of labour-intensive steps, and exert strong selection pressure. While the present work does not deliver a specific product beyond the characterised and evolved enzymes themselves, it establishes a modular framework that connects the assembly of genetic modules, the mining of sequences from large data sets, the evolution of enzymes through directed evolution and the evolution of synthetic yeast genome-enabled chassis, in order to select improved biocatalysis of displayed enzymes. The next step would be to connect this to downstream biosynthetic pathways in order to transform our platform into complete microbial yeast cell factories for the valorisation and upcycling of biopolymers.

### Study limitations

Several limitations of this study should be acknowledged. (i) The number and diversity of enzymes and SCRaMbLEd strains evaluated remain limited. (ii) Because the process is not yet coupled to cellular growth, pooled competition-based screening cannot be used to identify the best-performing candidates from large enzyme libraries and diverse SCRaMbLE populations. (iii) Depolymerisation was not integrated with a dedicated biosynthetic pathway or linked to biomass production to cover the whole range from the biopolymer waste stream to a product. Nevertheless, the framework established here provides a foundation for the systematic evaluation, optimisation and exploration of depolymerising enzymes, including their future integration into selection-based screening platforms and engineered bioproduction pathways.

## Materials and Methods

### Culture conditions

Unless stated otherwise, bacterial cultures were grown in 5 mL of LB medium (20 g/L tryptone, 20 g/L NaCl and 10 g/L yeast extract) in glass tubes at 37 °C overnight at 180–230 rpm on a 25 mm orbital shaker, supplemented with the respective antibiotic where necessary (ampicillin = 100 µg/mL, spectinomycin = 120 µg/mL). Yeast pre-cultures were inoculated from single colonies in 5 mL of YPD medium (20 g/L yeast extract, 40 g/L peptone, and with or without 0.64 g/L tryptophan) supplemented with 2% glucose if not stated differently, or synthetic complete medium (1.7 g/L yeast nitrogen base without amino acids and without ammonium sulfate, 5 g/L ammonium sulfate, 1.47 g/L SC triple drop-out -His, -Leu, -Ura; all components used are from Formedium) lacking the respective amino acid for plasmid maintenance with 2% glucose if not stated differently. Incubation was performed in glass tubes overnight at 30 °C at 180–230 rpm on a 25 mm orbital shaker.

### Strains, plasmids and oligonucleotides used and generated in this study

All yeast strains used and generated in this study are provided in Table S4. Details for plasmids are provided in Table S1 and as annotated GenBank files in Supporting Data. For this reason oligonucleotides for DNA assembly and sequence-validation purposes are not listed in this study; all other relevant oligonucleotides are listed with their relevant details in Table S5.

### Golden Gate assembly

Golden Gate cloning was performed using the standard procedure in 10-20 µL total volume or in 1 µL total volume using an acoustic dispenser (Echo 525 or Echo 650T) according to previously established protocols (Kobel et al., 2022; Kobel & Schindler, 2025). Briefly, 40 fmol (standard procedure) or 5 fmol (low-volume procedure) of each DNA part and the acceptor plasmid were combined with the respective type IIS restriction enzyme and T4 ligase (1 µL or 0.1 µL) in 1 × ligase buffer supplied, and ddH_2_O was added to reach the final reaction volume. The DNA assembly reaction was carried out using a thermal cycler and a cycling program [4 min 16 °C, 3 min 37 °C]_50_ Followed by a ligase inactivation step for 10 min at 50 °C and a restriction enzyme inactivation step for 10 min at 80 °C and storage at 8 °C until reaction mixture was transformed into RbCl-chemically competent *E. coli* cells (Hanahan, 1983). After heat shock for 30 seconds at 42 °C cells were recovered in 1 mL LB medium for 30 to 45 min with gentle agitation prior to plating on LB agar plates supplemented with the respective antibiotic and overnight incubation at 37 °C.

### Plasmid extraction and validation

Plasmid DNA was extracted using the BOMB protocol using carboxylated beads (Oberacker et al., 2019) according to the detailed protocol available at www.bomb.bio. Plasmids were tested by colony PCR or restriction pattern analysis using a suitable restriction enzyme prior to sequencing using either Sanger sequencing, whole plasmid sequencing or an established Nanopore-sequencing procedure from the group (Ramirez Rojas et al., 2024; Ramirez Rojas et al., 2025).

### Yeast transformation

Yeast strains were transformed as described previously (de Vries et al., 2024) according to a modified protocol described by Gietz and Woods (Gietz & Woods, 2002). Briefly, yeast strains were inoculated into 5 mL liquid medium and incubated overnight at 30 °C in a roller drum. Overnight cultures were then reinoculated into 20 mL liquid medium to OD_600_ = 0.1 and incubated in a roller drum to a target OD_600_ = 0.5. Cells were then centrifuged at 2000 × *g* for 5 min, washed with 10 mL sterile ddH_2_O and centrifuged again. Cell pellets were washed with 10 mL 0.1 M LiOAc and centrifuged at 2000 × *g* for 5 min. The cell pellet was resuspended in 400 µL 0.1 M LiOAc. 50 μL cells were mixed with up to 24 μL DNA, 90 μL 1 M LiOAc, topped up with ddH_2_O (depending on DNA volume: DNA + ddH_2_O = 24 µL), 10 μL 10 mg/mL salmon sperm carrier DNA (prior to each use boiled for 15 min at 100 °C and directly transferred to ice to obtain single-stranded DNA), and 600 μL 50% PEG 3350. Transformation mixtures were briefly mixed by vortexing and incubated at 30 °C for 30 min before the addition of 100 μL DMSO, followed by inversion or vortexing and heat shock at 42 °C for 15 min. Cells were pelleted at 1000 × *g* for 1 min, resuspended in 250 μL 5 mM CaCl_2_, incubated at room temperature for at least 10 min, and plated onto the respective selective medium agar plates.

### Construction of CRISPR/Cas sgRNA plasmids and repair DNA

As previously described (Lindeboom et al., 2024) sgRNA plasmids were prepared according to the protocol provided by the Ellis Lab (https://benchling.com/pub/ellis-crispr-tools) (Awan et al., 2017). Briefly, the generation of sgRNA plasmids was accomplished by insertion of annealed oligos into the destination plasmid pWS082 via Golden Gate cloning. Guide RNAs were designed using the online tool CHOPCHOP (Labun et al., 2019). Repair templates were generated using overlap extension PCR: 5′ and 3′ DNA segments flanking the area to be edited were amplified from genomic DNA using the corresponding designed oligonucleotides (Table S5). The resulting fragments contain an overlap with matching T_m_ for the 5′ amplicon forward and 3′ amplicon reverse primer to fuse both fragments in a subsequent PCR. 1 μL of each of the 5′ and 3′ amplicons was used in the second PCR as a template without purification, resulting in the repair template. The repair templates for CRISPR/Cas9 contained at least 300 bp for homologous recombination in yeast at both ends. Prior to yeast transformation, the sgRNA plasmids were linearized for 1 h by digestion with EcoRV.

### CRISPR/Cas9-mediated gene deletion in yeast

CRISPR/Cas9 was used to generate SLy0475. As previously described (Lindeboom et al., 2024) the preparation of sgRNA plasmids was done according to the protocol provided by the Ellis Lab (https://benchling.com/pub/ellis-crispr-tools) (Awan et al., 2017). Briefly, linear fragments encoding Cas9, sgRNA(s), and repair template(s) were cotransformed into yeast strains using the aforementioned yeast transformation method. Deletion was subsequently confirmed using colony PCR and Sanger sequencing. To prepare yeast cells for the colony PCR a colony was transferred into 40 µL of sterile water. 20 µL of this solution was mixed with 20 µL of 40 mM NaOH in a PCR reaction tube and boiled for 15 min at 100 °C in a PCR cycler. 1 µL of boiled cells was used as template in the colony PCR using a OneTaq polymerase with the respective PCR conditions in a total reaction volume of 10 µL. The remaining 20 µL of yeast cells were stored at 4 °C and used to recover the strain in case the confirmation was positive.

### ABTS colorimetric assay

To establish the assay, single colonies of yeast were inoculated and grown for 24 h at 30 °C in 96-well microtiter plates with 50 µL of SC-Leu medium supplemented with 2% raffinose and sealed. After 24 h each well was supplemented with 160 µL of expression medium (YPD, 2% galactose, 2 mM CuSO_4_, 60 mM KH_2_PO_4_ pH 6.0) and incubated at 30 °C to test three time points for optimal expression to measure laccase activity (24 h, 48 h and 72 h). Each time point was subjected to the ABTS assay to detect laccase activity. For the ABTS assay 20 µL of yeast culture with cells were mixed with 180 µL of activity buffer (100 mM sodium acetate pH 5.0, 2 mM ABTS (Sigma-Aldrich; A1888)). The oxidation of ABTS was measured at 418 nm in a plate reader (SPECTROstar^®^ Omega, BMG LABTECH). From this, 24 h was determined as the optimal time point at which laccase activity was detected.

### Directed evolution of laccase sequence through error-prone PCR

To obtain an average of 1-3 mutations per construct, different ep-PCR conditions were tested to determine the optimal conditions using Taq polymerase (Sigma-Aldrich; D1806). Three libraries of sLac_Amyco_ with varying mutation rates were generated by adjusting the concentration of MnCl₂ (0.01 mM, 0.1 mM, and 0.2 mM) during ep-PCR. The plasmid backbone (pSL1024) and the sLac_Amyco_ libraries were purified, DpnI digested and transformed into *S. cerevisiae* to assemble plasmids via *in vivo* homologous recombination, using the standard transformation procedure stated above. To determine suitable ep-PCR conditions for directed evolution, six transformation reactions were performed. In the three experimental conditions, 100 ng of amplified backbone plasmid was co-transformed with 300 ng of amplicon library generated in the presence of 0.01, 0.1, or 0.2 mM MnCl₂. As controls, 100 ng of amplified backbone plasmid was co-transformed with 300 ng of the sLac_Amyco_ amplicon (positive control), 100 ng of amplified backbone plasmid was transformed without PCR amplicon (negative control), and 100 ng of circular plasmid was transformed as a transformation control. The library with 0.2 mM MnCl_2_ resulted in an excessively high mutation rate with loss of function for the tested candidates and was therefore omitted from further analysis. Selected candidates were characterized using the ABTS assay. Directed evolution selected candidates exhibiting laccase activity in an initial screen underwent two validation rounds. First, 3 to 5 biological replicates of each candidate were tested with the ABTS assay. Second, plasmids were extracted from the selected candidates, and retransformed into the indicated *S. cerevisiae* strain background and the laccase activity assay was repeated to ensure that the activity was caused by the plasmid-encoded sequence and not by the strain background (e.g., random mutation). For selection in case of SCRaMbLEd yeast strains see section “*SCRaMbLE and selection of candidates*”. Plasmids from candidates with observed improved sLac_Amyco_ activity relative to the parental laccase version were subjected to Sanger sequencing to determine the underlying mutations.

### Functional annotation of enzymes in prokaryotic databases

High-throughput protein alignment was performed using the DIAMOND v.2.1.8 (Buchfink et al., 2015) algorithm to identify sequence similarities (at least 25% identity) between protein sequences in different metagenomic and genomic databases (GTDB r207, GMGB, GEM and Ocean (Rodriguez Del Rio et al., 2024)) with high-quality functional annotations available in KEGG (Kyoto Encyclopedia of Genes and Genomes, v.103.0) (Kanehisa & Goto, 2000). Each metagenomic sequence was assigned a KO code based on the best match obtained from the DIAMOND alignment against the KEGG database. An SQLite database was constructed to organize the annotated metagenomic sequences along with their corresponding KO codes. This database served as a comprehensive resource for subsequent analyses and queries and will be made accessible upon publication.

### Sequence and structural analysis of divergent laccases from databases

Conserved amino acid motifs along the protein sequences were identified with MEME v.5.4.1 (Bailey et al., 2015). Subsequently, structural prediction of the sequences with conserved amino acid motifs was carried out using AlphaFold2 server (Jumper et al., 2021). A comparison of the predicted sequences to the reference protein structures was done with TMALIGN (Zhang & Skolnick, 2005) and Foldseek (van Kempen et al., 2024). The metrics used to compare the degree of similarity between the structures were TMSCORE and RMSD. The metabolic potential assessment of genomes and metagenomes of putative laccases was done with MEBS v.1.2 (RRID:SCR_015708) (De Anda et al., 2017). MEBS does not provide the Markov models of lignin metabolism Pfams. For this reason, the custom Markov models files were generated based on literature of the respective biological functions (found in Table S6 and downloaded from InterPro (Hunter et al., 2009). In addition, the potential of the protein to be secreted was predicted using SignalP-6.0 (Teufel et al., 2022).

### Phylogenetic analysis of laccases

Two groups of orthologous laccase proteins were analysed, and phylogenetic trees were constructed to resolve their evolutionary relationships. Alignment and trimming were performed in Geneious Prime v2025.0.2 and MAFFT v7.490 (Katoh et al., 2002) (parameters: scoring matrix-BLOSUM 62). Poorly aligned sequences were manually removed with Geneious Prime v2025.0.2 and re-aligned with MAFFT v7.490. Alignment columns comprising >50% gaps were removed. Maximum-likelihood phylogenetic trees were constructed with IQTree v2.0.7 (Nguyen et al., 2015). The best model generated was selected according to the Bayesian information criterion (BIC). The two trees were constructed as follows: (1:K00421) LG+F+I+R5 with 1,000 bootstraps and (2:K10535) WAG+R8 model. The visual inspection of the evolutionary relationships was done with iTOL v.7.0 (Letunic & Bork, 2007).

### Sequence optimisation of mined sequences for DNA synthesis

All selected laccase sequences were codon harmonised for *S. cerevisiae* S288c using codonharmonizer (Claassens et al., 2017). Relevant type IIS restriction sites were eliminated using DNA Chisel (Zulkower & Rosser, 2020) and DNA fragments were synthesized by Twist Bioscience. The corresponding level 0 parts were cloned into the respective acceptor plasmid (pSL0103) and sequence validated. Level 0 plasmids were assembled into transcription units by Golden Gate assembly into the acceptor plasmid pSL1017. Information on the sequences mined will be made available upon publication in a peer-reviewed journal, and will not be disclosed in this preprint.

### Kraft-lignin depolymerisation analysis via HPLC-MS

Single colonies with the respective plasmid (pSL1179, pSL1160 and pSL1017), were grown in 5 mL SC-Leu with 2% raffinose overnight at 30 °C. The next day, an expression culture was inoculated in a 250 mL flask with a total volume of 50 mL, starting at OD_600_ = 0.01, and grown for 24 h at 30 °C. For HPLC-MS analysis a 2 mL reaction was prepared with a final OD_600_ of 10, supplemented with 0.1% kraft-lignin (w/v) and 2 mM ABTS. Samples were generated as triplicates and incubated at room temperature. Samples were collected at 0, 2, 4, 6, 24, 48, and 72 h and immediately stored at -80 °C until extraction.

Untargeted metabolic profiling was performed using HRES-LC-MS/MS. Chromatographic separation was carried out on a Vanquish LC system (Thermo Scientific) applying two complementary chromatographic methods: Compounds of moderate polarity were chromatographically separated using a Kinetex EVO C18 column (150 × 2.1 mm, 1.7 μm particle size, Phenomenex) coupled to a matching guard column (20 × 2.1 mm, 3 μm particle size, Phenomenex). The column temperature was maintained at 40 °C, and the mobile phase flow rate was set to 0.250 mL/min. A gradient elution method was employed using 0.1% formic acid in water (phase A) and 0.1% formic acid in methanol (phase B), with a total run time of 15 minutes. The mobile phase flow was structured in the following steps and linear gradients: 0 – 1 min constant at 30% B; 1 – 8 min from 30% to 100% B; 8 – 10 min constant at 100% B; 10 – 10.10 min from 100 to 30% B; 10.10 – 15 min constant at 30% B. Compounds of high polarity were chromatographically separated using a SeQuant ZIC-pHILIC column (150 × 2.1 mm, 5 μm particle size, peek coated, Merck) connected to a guard column of similar specificity (20 × 2.1 mm, 5 μm particle size, Phenomoenex) a constant flow rate of 0.1 mL/min with mobile phase A comprising 10 mM ammonium acetate in water, pH 9, supplemented with medronic acid to a final concentration of 5 μM (A) and 10 mM ammonium acetate in 90:10 acetonitrile to water, pH 9 (B) at 40 °C. The mobile phase profile consisted of the following steps and linear gradients: 0 – 1 min constant at 75% B; 1 – 6 min from 75 to 40% B; 6 to 9 min constant at 40% B; 9 – 9.1 min from 40 to 75% B; 9.1 to 20 min constant at 75% B. The injection volume was 5 µL for both types of chromatography.

Mass spectrometric detection was performed using an Orbitrap ID-X mass spectrometer (Thermo Scientific) equipped with a high-temperature electrospray ionization (HESI) source either in positive ion or negative ion mode (separate injections). The instrument was operated at a static spray voltage of 3500 V (positive mode) or 2500 V (negative mode), with the following parameters: sheath gas: 50 arbitrary units (Arb); auxiliary gas: 10 Arb; ion transfer tube temperature: 325 °C; vaporizer temperature: 350 °C. Data-dependent MS² acquisition was performed at an Orbitrap mass resolution of 60,000 with quadrupole isolation in the m/z range of 100-1000, followed by high-energy collision dissociation (HCD) at a relative collision energy of 30%. Fragment ions were detected using the Orbitrap mass analyzer at a mass resolution of 30,000. To increase spectral coverage, dynamic exclusion was applied with an exclusion time of 2.5 seconds and a mass tolerance of 10 ppm.

Mass spectrometer raw data were aligned and peak-integrated using MS-DIAL (5.5) with the following parameters: MS1 tolerance (0.01 Da), MS2 tolerance (0.025 Da), retention time tolerance (0.1 min), peak count filter (15%), mass slice width (0.05 Da), and minimum peak height (10,000). The resulting exported file containing the aligned, integrated peaks, and list of MS1 and MS2 spectral features, was used as input in Sirius 6.3.0 to run the built-in tools: Sirius, CSI:FingerID, and CANOPUS. The resolution was set to 5 ppm, and selected for the Orbitrap instrument. Molecular formula generation was configured using a combination of *de novo* (for m/z values below 400) and bottom-up approaches. Features above 850 m/z were included in the computation. Search structural databases included GNPS, COCONUT, PlantCyc, SuperNatural and PubChem. Two output files from Sirius were used in the analysis script: (1) the list of top molecular structures for each identified formula, and (2) the list of ClassyFire ontology annotations for each feature according to the highest-confidence structure. The MS-DIAL and Sirius outputs were then combined. Features were retained for annotation if they passed blank filtering (sample-to-blank intensity ratio ≥ 10, i.e. blank signal ≤ 0.1 of sample signal). Putative structural annotations were then filtered to retain only high-confidence hits, with a minimum SIRIUS confidence score of 0.9 and a minimum class probability of 0.9. Normalization was done by total ion current (TIC). One-way ANOVA was used to identify compounds with statistically significant changes in signal intensity across time and samples. For compounds selected as lignin-degradation–associated, z-scores were calculated across samples and visualised as a heatmap to represent relative abundance patterns.

### Cre-mediated lethality assay of synthetic yeast strains

The Cre recombinase expression plasmid (pSL0217 or pSCW11-CreEBD (synXI and synXII)) was transformed into yeast cells as well as the pRSII413 as a control plasmid for all strains, except synXI and synXII which were transformed with the control plasmid pRSII416 due to an existing genomically integrated HIS3 in the strains. Each experiment was performed in three biological replicates. Yeast pre-cultures were inoculated from single colonies in 5 mL SC-His or SC-Ura, 2% glucose and incubated at 30 °C. The next day, 5 mL of the respective SC medium were inoculated to a starting OD_600_ of 0.1. Three treatments were prepared for each strain as follows: i) strain + Cre-plasmid with *β*-estradiol, ii) strain + Cre-plasmid without *β*-estradiol, and iii) strain + pRSII413 or pRSII416. For Cre recombinase induction with *β*-estradiol (Sigma-Aldrich, E2758) a final concentration of 1 μM was used. Samples were taken at 0, 2, 4, 6 and 24 h. For each time point a 10-fold dilution series in a 96-well microtiter plate was generated starting from 200 μL culture. The dilution series was plated onto single-well YPD agar plates using the ROTOR HDA from Singer Instruments. A custom 7 × 7 program was used, with the source plate revisited at the start of each row of the printed patch. Plates were incubated for 48 h at 30 °C. CFUs were detected using ImageJ (Schindelin et al., 2012) with the Count-On-It plugin (Dodge & Ludington, 2023) for automated detection of CFUs. Detected CFUs were manually verified with the 10^-3^ dilution. The means of CFUs for the three biological replicates were calculated and the CFUs induced and non-induced were normalised by the mean CFUs count at t = 0 h of the non-induced treatment. The lethality factor was calculated by the ratio of CFUs induced divided by CFUs in the uninduced treatment.

### Plate reader growth curve kinetics of yeast

Growth curve analysis was performed using a SPECTROstar^®^ Omega, BMG LABTECH plate reader in 96-well flat bottom plates (Greiner; 655180) with 150 µL of medium inoculated to a starting OD_595_ of 0.05. Plates were sealed with TopSeal-A PLUS (Perkin Elmer; 6050185). The following kinetic parameters were run with a varying number of cycles (approx. 24 h at 30 °C: 2:00 min orbital shaking 2:00 min linear shaking followed by OD measurement at 595 nm. Data was analysed and visualised using the R package GrowthCurver (Sprouffske & Wagner, 2016) and custom scripts.

### SCRaMbLE and selection of candidates

Synthetic yeast strains with Cre recombinase expression plasmid (pSL0217 or pSCW11-CreEBD) were transformed with pSL1160 and after validation streaked on SC-Leu-His or SC-Leu-Ura plates to obtain single colonies. Pre-cultures from single colonies were cultivated in 5 mL of the respective SC-medium supplemented with 2% glucose at 30 °C overnight. The next day, 5 mL cultures were inoculated with a starting OD_600_ of 0.1 in the same medium with and without 1 µM *β*-estradiol and cultivated for 6 h. 200 µL of 10^-1^ and 10^-2^ dilution were plated on SC-Leu plates and incubated at 30 °C for 48 h. Subsequently laccase activity was determined as described before. Relevant candidates were analysed for their laccase activity, normalised by cell count (Martinic Cezar et al., 2025). Briefly, three single colonies of each candidate strain expressing the sLac_Sspir_ (pSL1160) as well as the control strains having an empty plasmid (pRSII413) were picked as replicates in 96-well microtiter plates containing 150 μL of SC-Leu supplemented with 2% raffinose and incubated at 30 °C for 24 h. The cultures were used to inoculate a 20 mL expression culture in 100 mL Erlenmeyer flasks and grown for 24 h in an orbital shaker with 25 mm diameter at 230 rpm. OD_600_ was measured and adjusted to a final OD_600_ ∼80 in one mL. The correct cell density was determined by a dilution series.

Laccase activity of 50 µL of the cell suspension was measured in a 1.5 mL reaction tube by adding 450 µL assay buffer (100 mM sodium acetate buffer pH 5.0, 2 mM ABTS). To measure laccase activity at 418 nm for T0 and T1, cells in the reaction tube were centrifuged at 2000 × *g* for 2 min and 200 μL of the supernatant were taken for the measurement. The remaining cells and reaction buffer were gently shaken and incubated in the dark in the presence of oxygen. After ∼12 h, cells were concentrated again at 2000 × *g* for 2 min and 200 μL were taken to measure T1 at 418 nm. For each biological replicate, the difference in absorbance at 418 nm was calculated and normalised to the cell counts as follows:

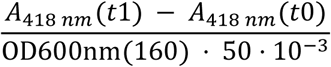

Formula explanation: 160 is the dilution factor (5 µL cells in 795 µL) and 50 × 10^-3^ is the number of cells added for activity measurement in mL (here 50 µL) multiplied by the dilution factor.

### High-molecular weight gDNA extraction for long-read sequencing

Genomic DNA was obtained as described previously (Lindeboom et al., 2024) using the NucleoBond HMW DNA kit (Macherey-Nagel; 740160) according to the manufacturer’s guidelines using lyticase (Sigma-Aldrich; L4025) for cell lysis (25 μL of 10,000 U/mL stock in Potassium Phosphate Buffer (10 mM K_2_HPO_4_, 10 mM KH_2_PO_4_, pH 7.0 in 50% glycerol)) for 1 h at 37 °C in 1.5 mL of Y1 buffer (1 M sorbitol, 100 mM EDTA pH 8.0, 14 mM *β*-mercaptoethanol). DNA quality and concentration were assessed via gel electrophoresis, NanoDrop spectrophotometer, and the Qubit 3 fluorometer using dsDNA BR reagents.

### Nanopore sequencing and data analysis

Library preparation was performed using the Ligation Sequencing kit SQK-NBD114.24 (Oxford Nanopore Technologies) with Native Barcoding kits EXP-NBD104 and EXP-NBD114 for multiplexing. Sequencing was performed on a MinION Mk1B device using MinION Flow Cells FLO-MIN114 (R10). Kits were used according to the manufacturer guidelines, except the input DNA was increased 5-fold to match the molarity expected in the protocol as no DNA shearing was applied. In previous testing we observed our fragments to be predominantly around 50 kb while the protocol is based on 10 kb DNA fragment length input. Sequencing experiments were run for 72 hours. Basecalling was performed using Dorado (version: 1.3.2, Oxford Nanopore Technologies) and sequencing data will be made accessible upon publication under BioProject ID: PRJNA1532932. Reads were mapped to the *S. cerevisiae* S288c reference (PRJNA128) where the respective synthetic chromosome sequence replaced the native sequence using minimap2 (version 2.17-r941) (Li, 2018). Flye (2.9.6-b1802) (Kolmogorov et al., 2019) was used to perform *de novo* assembly of chromosomes of individual sequencing experiments. The assembly quality was determined using Quast (version 5.3.0) (Gurevich et al., 2013), followed by correction based on the reference to remove potential mis-assemblies using RagTag (version 2.1.0) (Alonge et al., 2019). Variant calling was performed with svim-asm (1.0.3) (Heller & Vingron, 2019) and with sniffles (v 2.3.3) (Sedlazeck et al., 2018) using the mapped reads. Detected variants were manually curated based on mapped reads using the IGV web server (Robinson et al., 2011). Additionally, dot plots to compare reference genomes with the SCRaMbLEd candidates were generated using MUMmer (version 3.5) (Delcher et al., 1999) for rapid assessment of SVs.

### Statistical analysis

One-way ANOVA statistical analyses were followed by pairwise post-hoc comparisons with Holm correction. Kruskal-Wallis tests were followed by Dunn’s post-hoc test. All analyses were performed in a Jupyter notebook using Python v3.13. The test used is indicated in the corresponding figure legend unless stated otherwise.

## Supporting information

● Supporting Information: Supporting Figures, Supporting Tables, and Supporting References

● Supporting Data: GenBank files of all relevant plasmids and DNA sequences

## Material availability statement

All data and materials generated or analysed during this study are included in the article and its Supplementary Information, or will be made available upon the article’s publication in a peer-reviewed journal. Long-read sequencing data are deposited under BioProject PRJNA1532932 and will be available upon publication. The plasmids and strains generated during this study are available from the corresponding author upon request, except for the synthetic yeast strains. These were kindly provided by Jef D. Boeke (NYU, New York, USA) and should be requested directly from his laboratory.

## Acknowledgments

We would like to acknowledge the contributions of all current and former members of the Schindler Lab at the Max Planck Institute for Terrestrial Microbiology and the Center for Molecular Biology at Heidelberg University (ZMBH). In particular, we would like to thank Alexa Weikert for her support with strain construction and SCRaMbLE experiments, and Sally Jones for proofreading the manuscript. We would also like to thank all the other team members from our collaborating institutions for their input, discussions and support. We are very grateful to Jef D. Boeke (NYU, New York, USA) for sharing the synthetic yeast strains. We thank Tom Ellis (Imperial College London, UK) for sharing the CRISPR plasmids pWS082 and pWS158. This work was funded by the Max Planck Society as part of the MaxGENESYS project (DS), as well as through core funding of the Schindler Lab at ZMBH. Additional support came from funding by the Carl Zeiss Foundation and the Center SynGen (DS), the European Union (NextGenerationEU) via the European Regional Development Fund (ERDF) from the state of Hesse for the project *’Biotechnological production of reactive peptides from waste streams as lead structures for drug development*’ (DS), and two grants from the Federal Ministry of Education, Research and Technology (BMBF) for the CaptureExpress (01DN23012) and TABASCO (031B1584) projects (DS/MMZ). MdCSO acknowledges an International Max Planck Research School (IMPRS) fellowship and a Research Travel Grant (RTG) from the Federation of European Microbiological Societies (FEMS) for performing directed evolution experiments and receiving training at EGR’s laboratory. EGR acknowledges funding from the Consejo Superior de Investigaciones Científicas (CSIC) under Grant I3-2024 PIE_2024ICT163.

## Author contributions

DS conceived and designed the study. MdCSO identified laccases and lignin degradation as the proof-of-concept scenario. MdCSO performed most of the experimental work. MdCSO and NP performed the metabolome measurements. MdCSO, JGI, JHC, EGR, VD, and DS performed the data analysis and interpretation. DS supervised the project and acquired funding. MdCSO and DS wrote the manuscript, with input from all authors. All authors reviewed and approved the final version.

## Declaration of interest

None to declare.

## Notes

### Competing Interest Statement

The authors have declared no competing interest.

