## Supporting Information for "Coupled enzyme discovery, evolution and synthetic yeast chassis adaptation for microbial biopolymer valorisation"

**Content:**

Supporting Figures S1-S14

Supporting Tables S1-S6

Supporting References

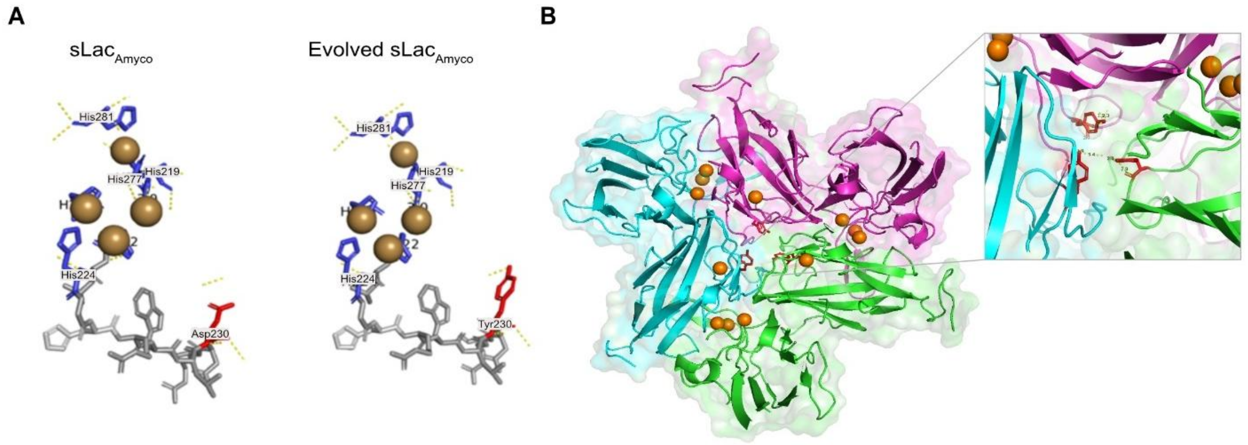


**Figure S1 | Mapping of the underlying D230Y mutation on the sLac$_{Amyco}$ protein structure. (A)** A close-up of D230Y, which increases the stability and observed activity of the evolved sLac$_{Amyco}$ relative to the parental protein sequence. **(B)** Mapping of D230Y onto the homotrimer of sLac$_{Amyco}$, with a close-up of the mutated residues highlighted in red. Each monomer is represented in pink, green and blue, respectively. Copper ions are shown as orange spheres. This model was generated using the PyMOL Molecular Graphics System (version 3.1.6.1, Schrödinger, LLC), based on the crystal structure of sLac$_{Amyco}$ (PDB: 3TA4).

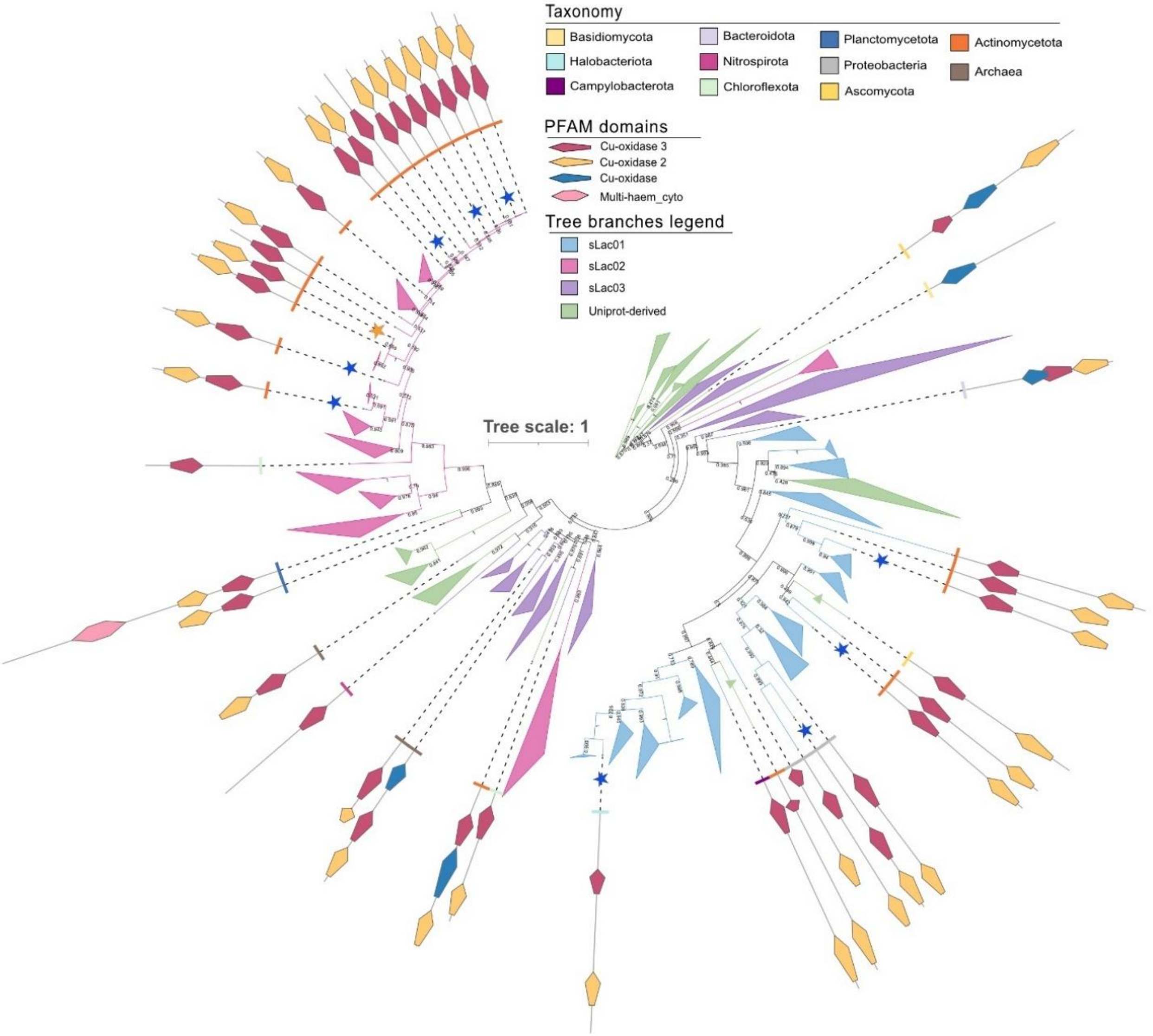


**Figure S2 | Phylogenetic tree of putative divergent small laccases.** This maximum-likelihood phylogeny of mined sequences was constructed using FastTree. The scale bar indicates substitutions per site. The sequences recovered in the mining approach form three subclades: sLac01 (blue), sLac02 (pink) and sLac03 (purple). The reference and reviewed sequences retrieved from UniProt for phylogenetic context are shown in green. Coloured wedges and tick marks around the tree indicate the taxonomic class of each sequence according to the legend. Selected putative divergent small laccases chosen for further experimental characterisation are marked with blue stars, and $sLac_{Amyco}$ is marked with a yellow star. PFAM domain architectures (Cu-oxidase 1, 2 and 3, as well as multi-heme cytochrome domains) are shown for representative sequences and are connected to their corresponding tips by dashed lines.

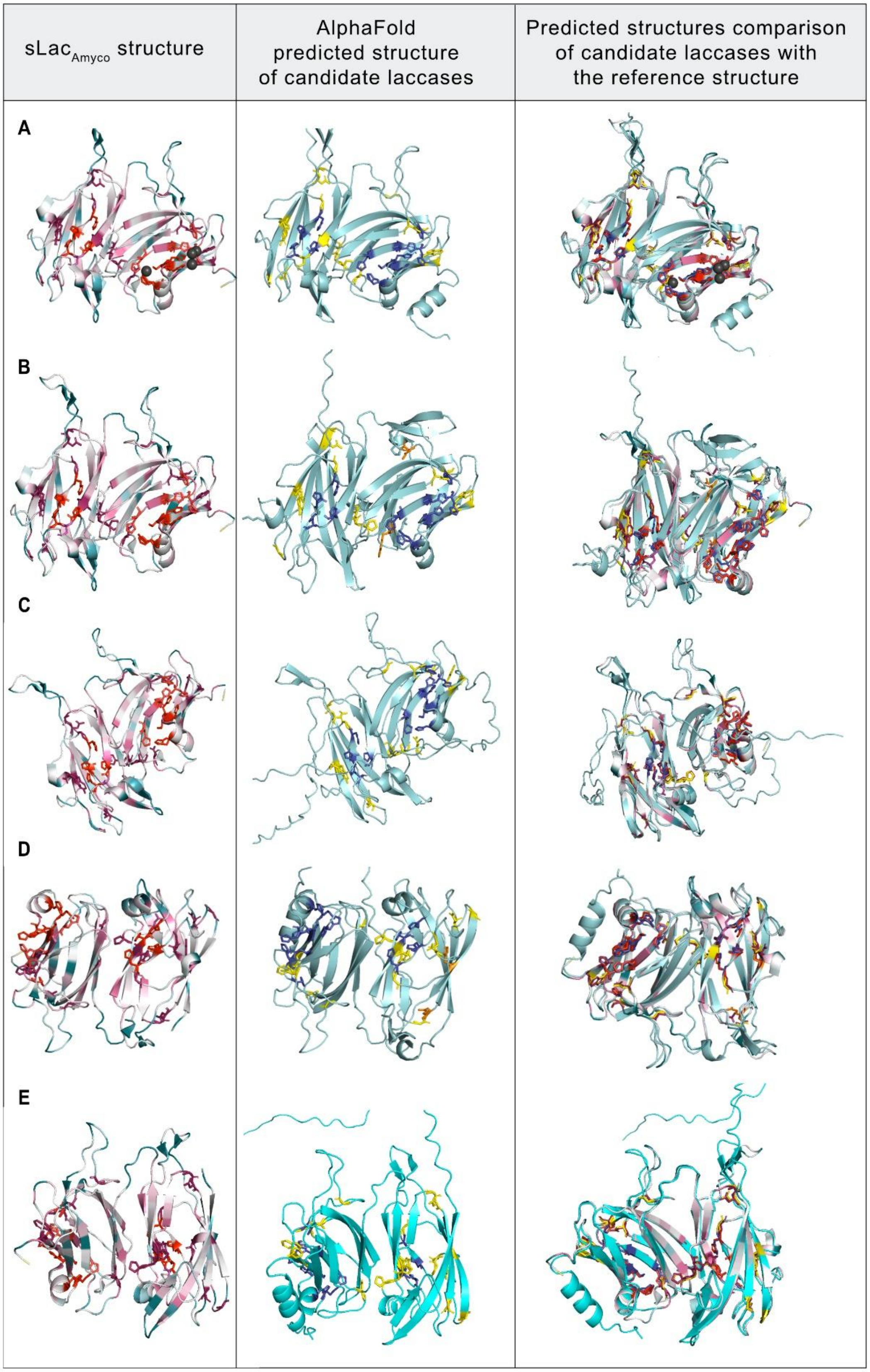


[Figure legend on following page]

**Figure S3 | Structural comparison of the predicted candidate small laccase structures.** From left to right: the AlphaFold-predicted structures of the candidate laccases; and the overlay of these two structures. **(A)** *A. xinjiangensis*, **(B)** *M.* sp. 016745425, **(C)** *S. spiralis*, **(D)** *P. flavigriseum*, and **(E)** *M. violae*. The models were generated using the PyMOL Molecular Graphics System (version 3.1.6.1, Schrödinger, Inc.). The level of conservation for $sLac_{Amyco}$ was determined using ConSurf. Dark pink represents the most conserved residues and red represents residues that are part of the active site and responsible for copper coordination. For the predicted structures of the candidate laccases, the most conserved residues compared with $sLac_{Amyco}$ are shown in yellow, the residues that form the active site are shown in blue, and the residues that differ from the conserved residues found in the sLacAmyco structure are shown in orange.

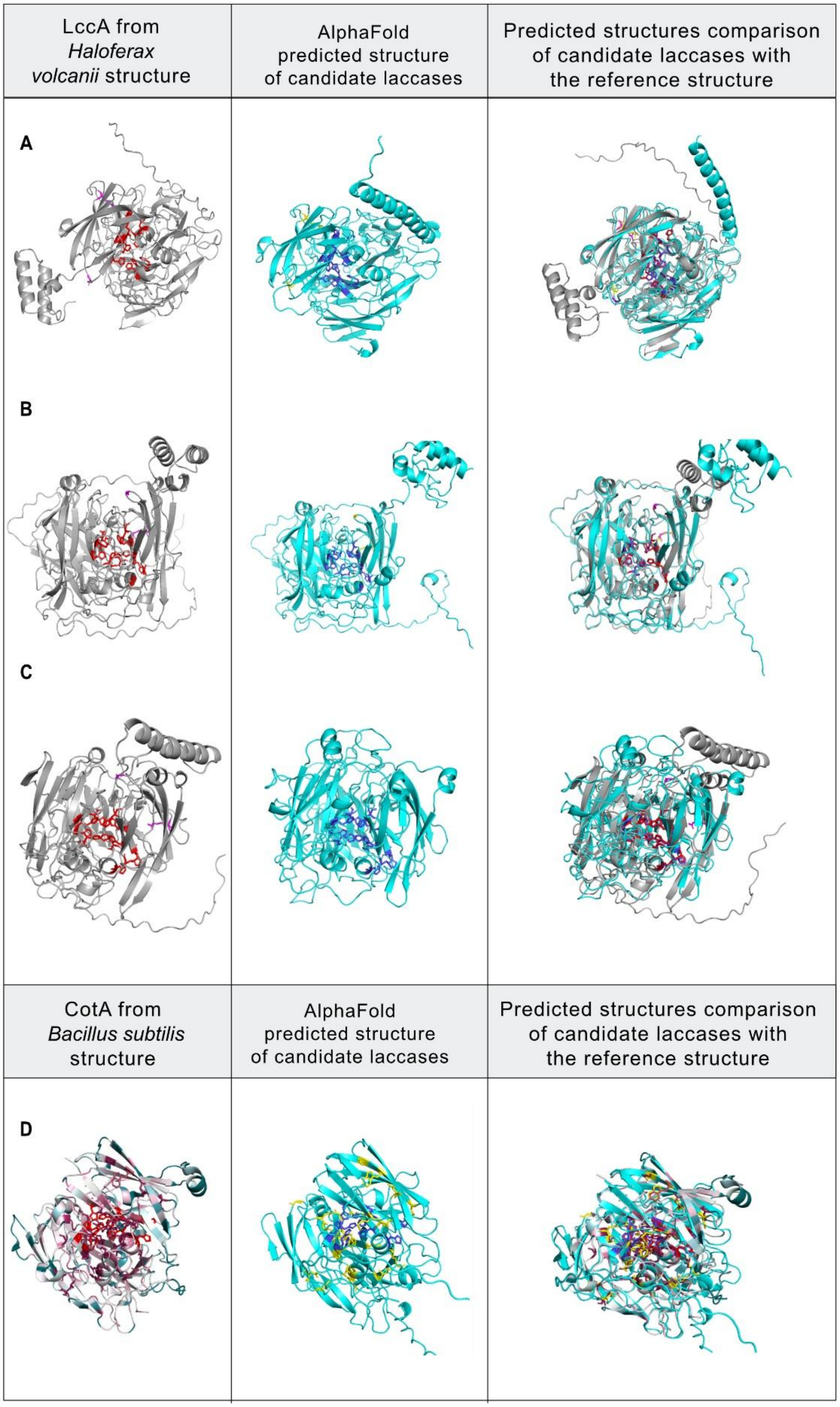


[Figure legend on following page]

**Figure S4 | Continuation of the comparison of the predicted structures for candidate small laccases.** From left to right: (A to C) LccA from *H. volcanii* (UniprotID: D4GPK6) and (D) CotA from *B. subtilis* (PDB: 7Y8C); the AlphaFold predicted structure of the candidate laccase; and the overlay of the two structures: **(A)** SIRX01 sp004563715, **(B)** *H. sedimenticola*, **(C)** *P. resinovorans*, and **(D)** *A. bryophytorum*. The models were generated using the PyMOL Molecular Graphics System (version 3.1.6.1, Schrödinger, Inc). The level of conservation for CotA from *B. subtilis* was determined using ConSurf. Pink represents the most conserved residues and red represents residues that are part of the active site and are also responsible for copper coordination. For the predicted structures of candidate laccases, the most conserved residues compared with the respective reference structure are shown in yellow, the residues that form the active site are shown in blue, and the residues that differ from the conserved residues found in the reference structure are highlighted in orange.

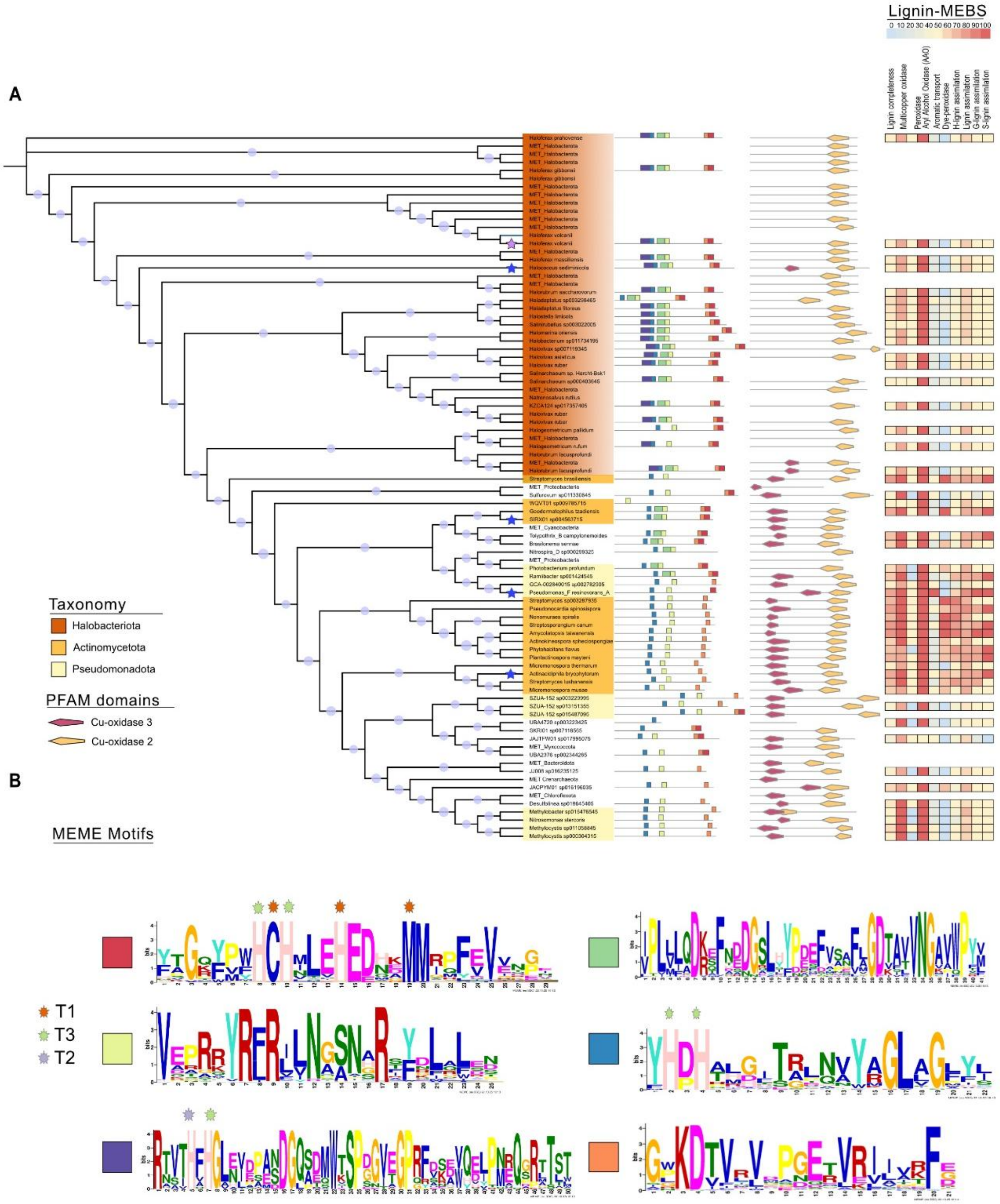


**Figure S5 | Phylogenetic tree of sLac01. (A)** Phylogenetic tree of sLac01 sequences that are at least 25% similar to the KEGG orthogroup K00421 of prokaryotic laccases. From left to right, the tree displays the following: (i) motif sequence conservation, (ii) conserved PFAM domains across species, and (iii) metabolic completeness for lignin degradation. This is shown using a set of enzymes involved in lignin degradation and assimilation, as determined by MEBS. Blue stars indicate the selected candidate laccases, while the purple star indicates the laccase from *H. volcanii*. **(B)** Detailed MEME motifs detected in the candidate laccases of sLac01. The residues responsible for forming copper clusters in laccases are highlighted in stars.

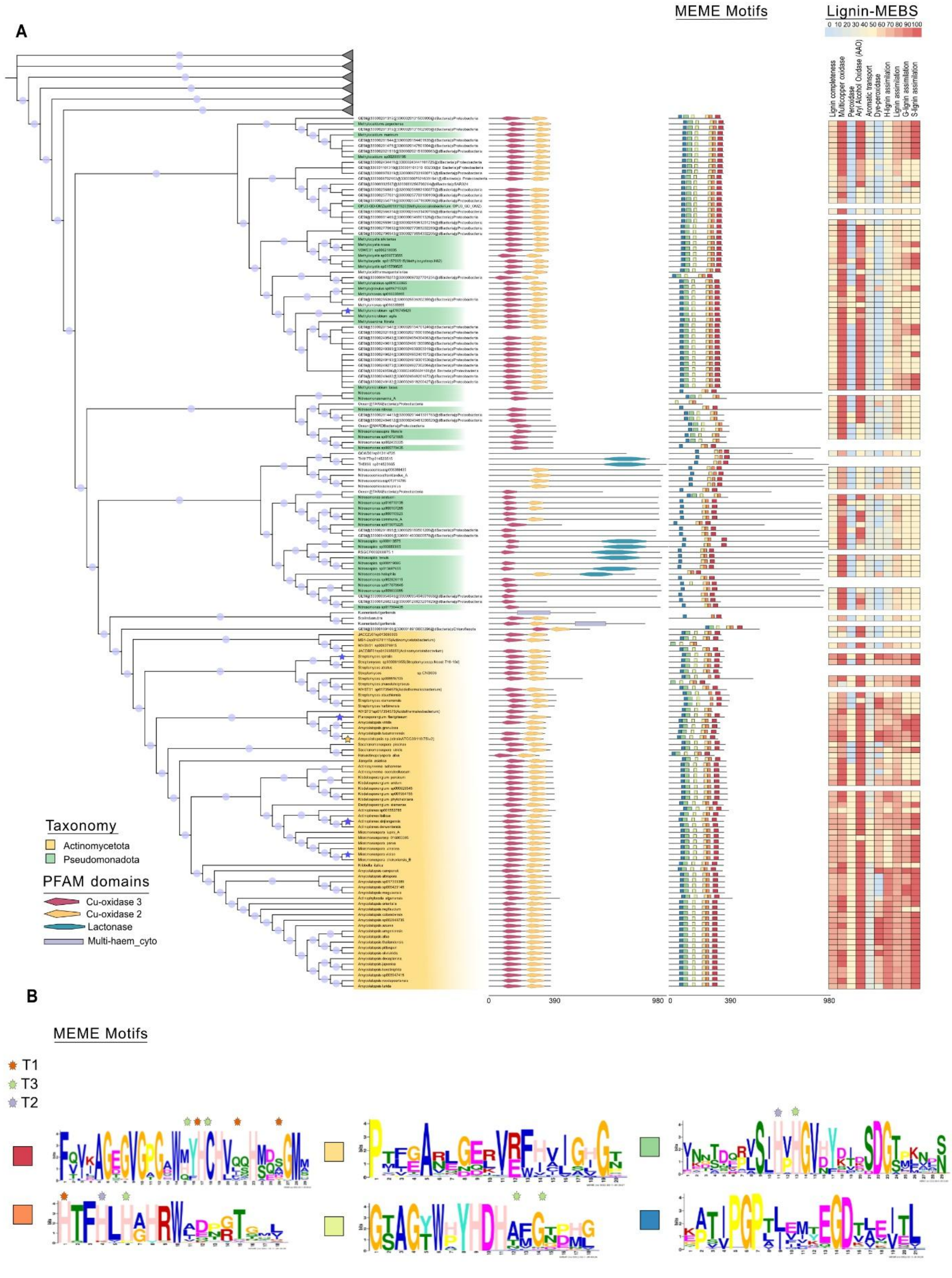


[Figure legend on following page]

**Figure S6 | Phylogenetic tree of sLac02. (A)** Phylogenetic tree showing sLac02 sequences that are at least 25% similar to the KEGG orthogroup K10535, which comprises hydroxylamine dehydrogenases. From left to right, the tree displays the following: (i) conserved PFAM domains across species; (ii) motif sequence conservation; and (iii) metabolic completeness for lignin degradation. This is shown for a set of enzymes involved in lignin degradation and assimilation using MEBS. Blue stars indicate the selected candidate laccases, and the yellow star indicates sLac$_{Amyco}$. **(B)** Detailed MEME motifs detected in the candidate laccases in sLac02. The residues responsible for forming copper clusters in laccases are highlighted with stars.

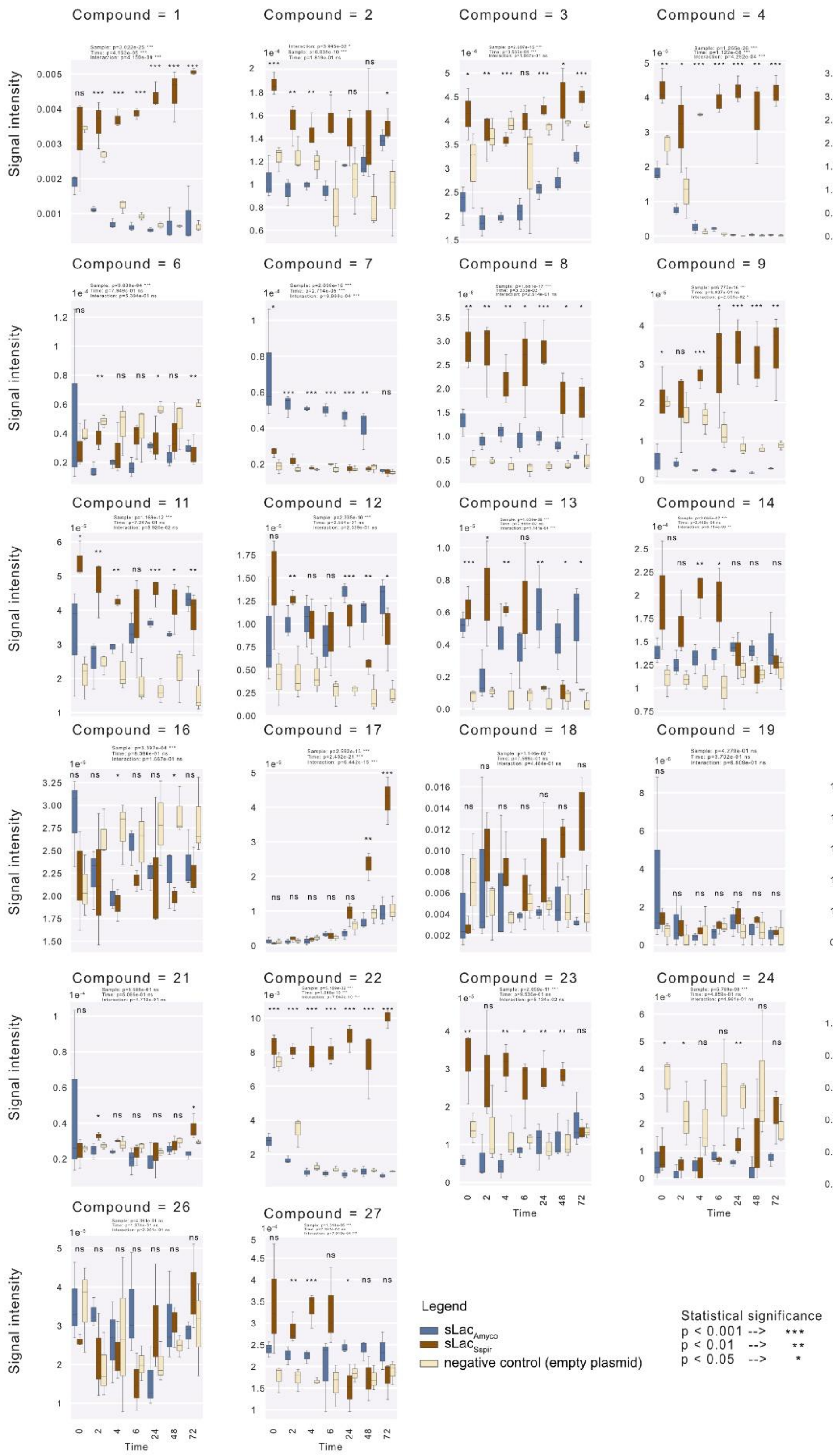


[Figure legend on following page]

**Figure S7 | Effect of treatment and incubation time on compound signal intensity.** The box plots show the signal intensity of the 27 compounds identified in this study (Table S3) across the three treatment conditions ($sLac_{Amyco}$, $sLac_{Sspir}$ and the negative control) at each of the seven sampling time points. A two-way ANOVA was performed for each compound with treatment and incubation time as fixed factors to assess their main and interactive effects on signal intensity. Statistical significance is indicated as follows: * $p < 0.05$, ** $p < 0.01$, *** $p < 0.001$, and ns: not significant; n = 3.

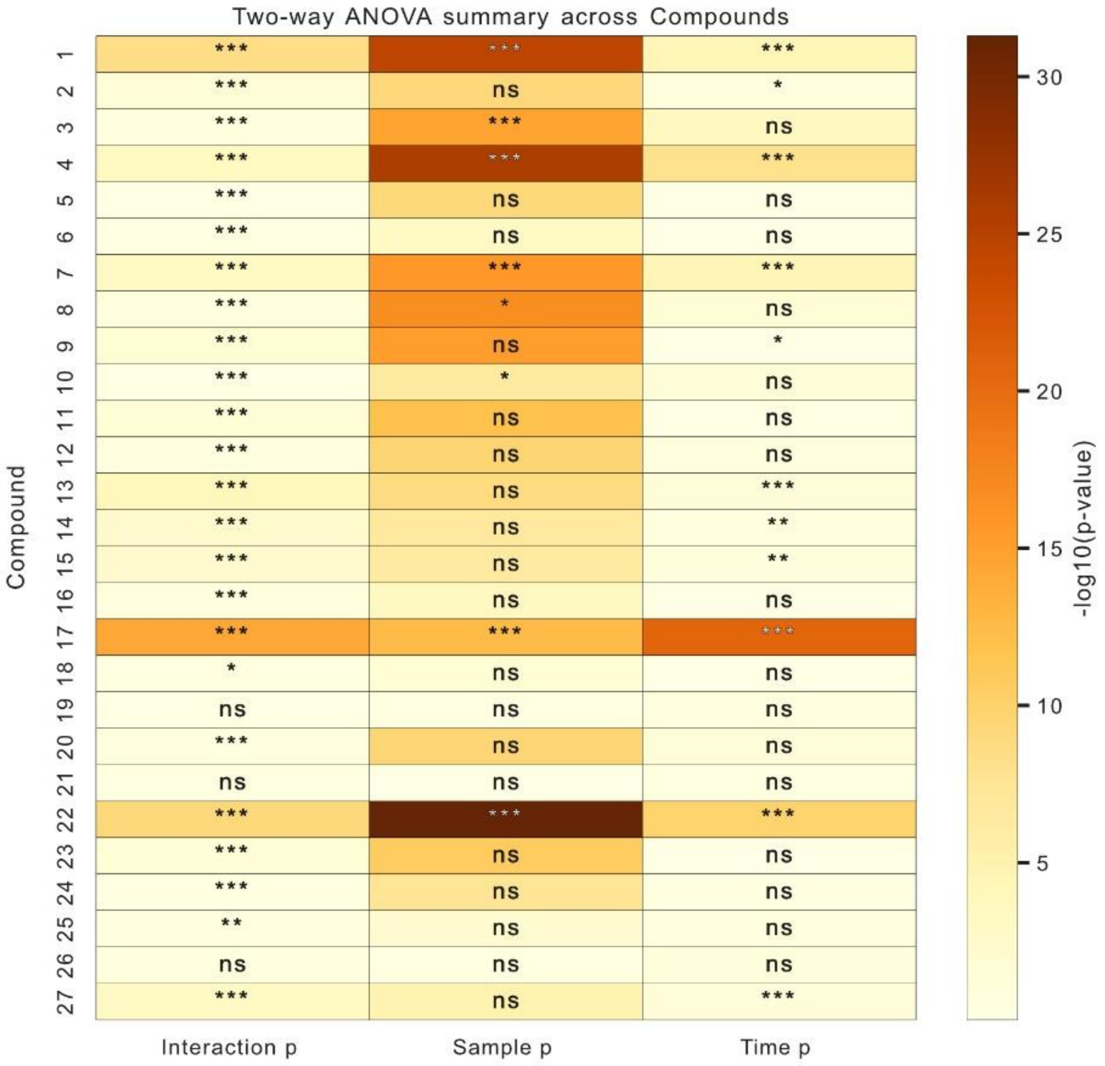


**Figure S8 | Summary of the results of the two-way ANOVA for the effects of treatment and incubation time on compound signal intensity.** Heatmap summarises the two-way ANOVA results for the 27 compounds identified in this study (Figure 3 and S7). The effects of treatment (sLac$_{Amyco}$ vs. sLac$_{Sspir}$ vs. the negative control), incubation time and their interaction (treatment × time) on signal intensity were tested. The colour scale represents the $-\log_{10}$ *p*-value for each factor, with darker shading indicating stronger statistical significance. Asterisks denote *p*-value thresholds (* $p < 0.05$, ** $p < 0.01$, *** $p < 0.001$, and ns: not significant).

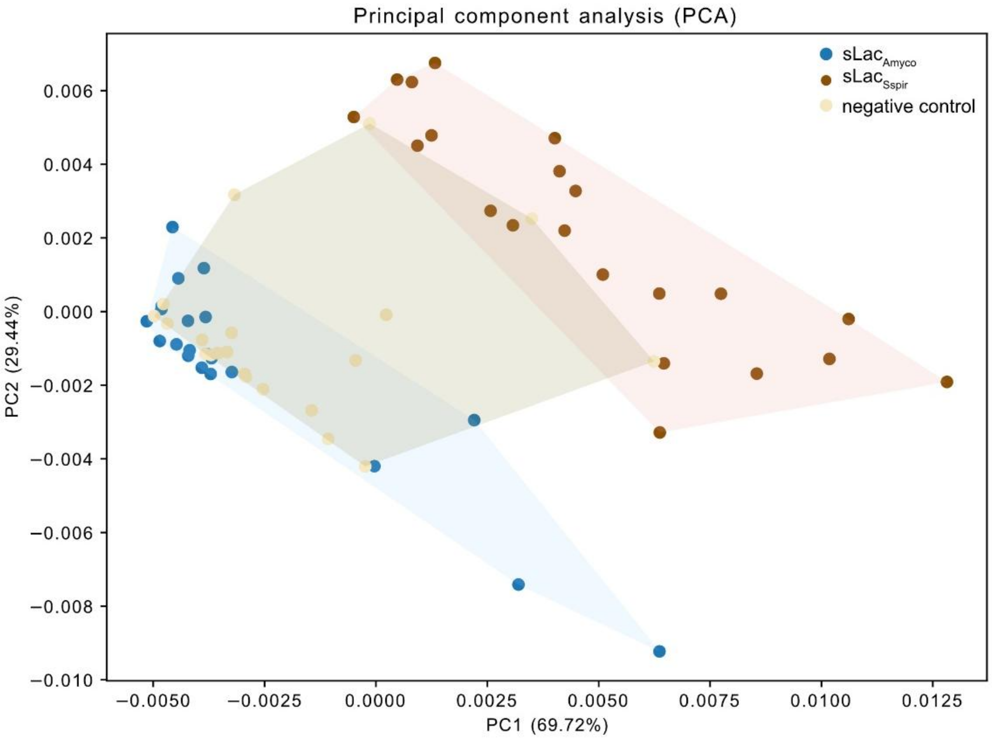


**Figure S9 | Principal component analysis (PCA) of compounds derived from lignin depolymerisation under different treatment conditions.** The PCA was performed on compounds detected following the treatment of Kraft-lignin with $sLac_{Amyco}$ and $sLac_{Sspir}$, as well as a negative control. Each point represents an individual sample, with shaded polygons denoting the convex hull for each treatment group. The data include three replicates per condition at each of seven time points (n = 21 per group). PC1 and PC2 explain 69.72% and 29.44% of the total variance, respectively.

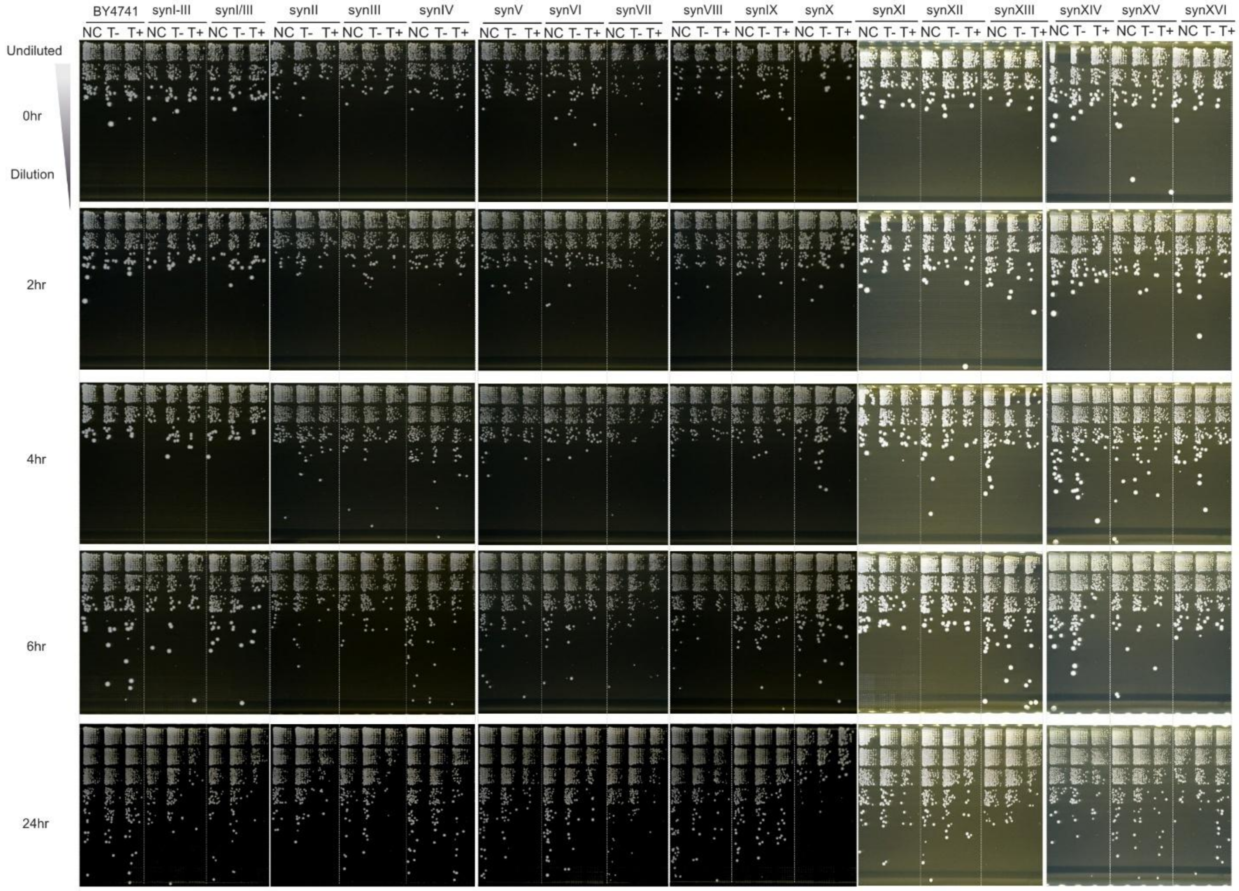


**Figure S10 | Lethality assay for the sixteen synthetic yeast strains.** Solid media dilution across time points for the sixteen synthetic yeast strains. NC: negative control (plasmid with auxotrophy marker for selection: pRSII413 or pRSII416). T: treatment condition (plasmid expressing Cre recombinase (pSL217 or pSCW11-CreEBD)). '-' refers to the absence of *β*-estradiol, while '+' refers to its presence.

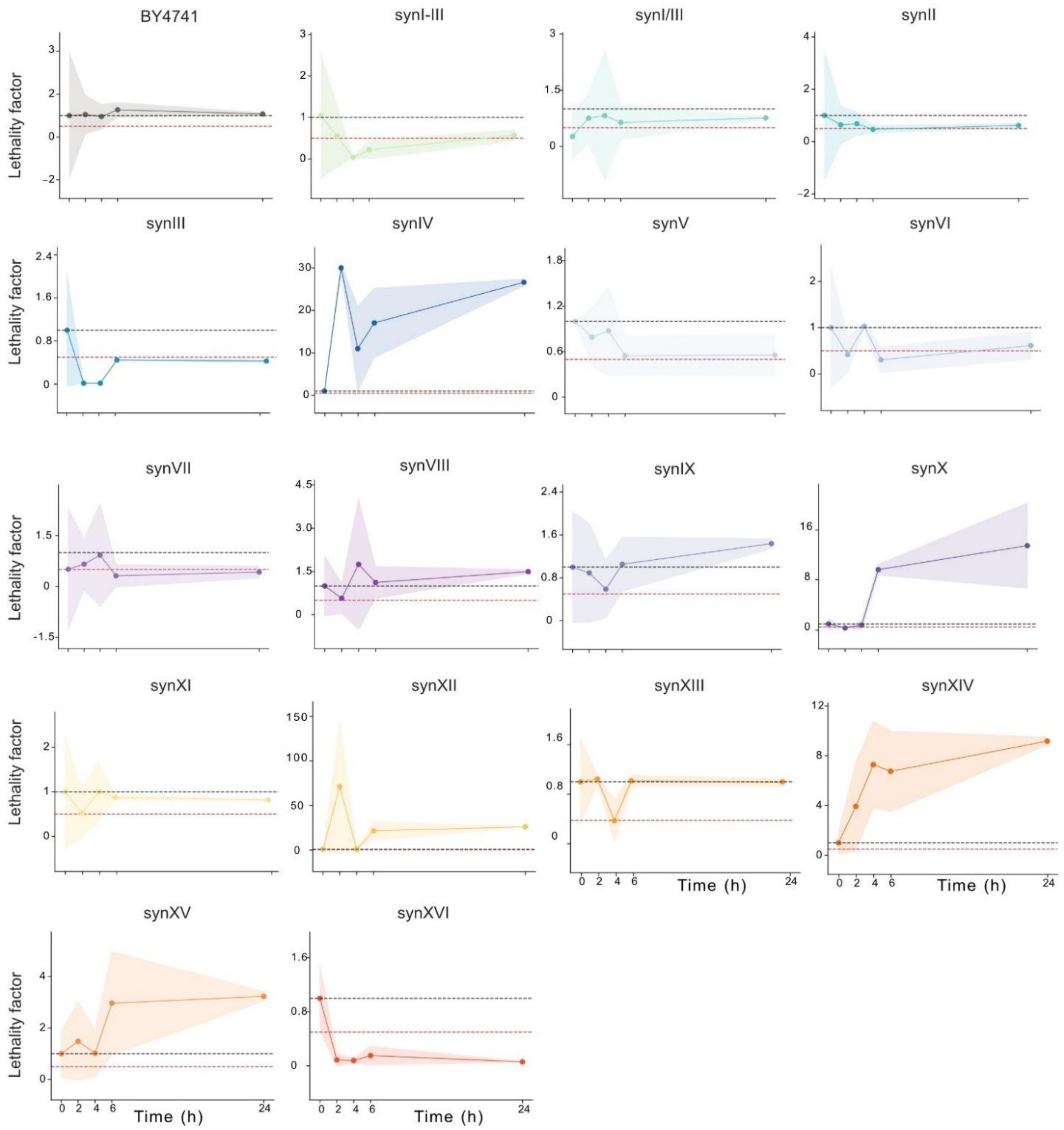


**Figure S11 | Lethality assay (Figure S10) visualized in plots for the sixteen synthetic yeast strains.** The non-continuous black line is set to 1 and the non-continuous red line to 0.5 over a 24-hour time course. The background indicates the standard deviation (n = 3).

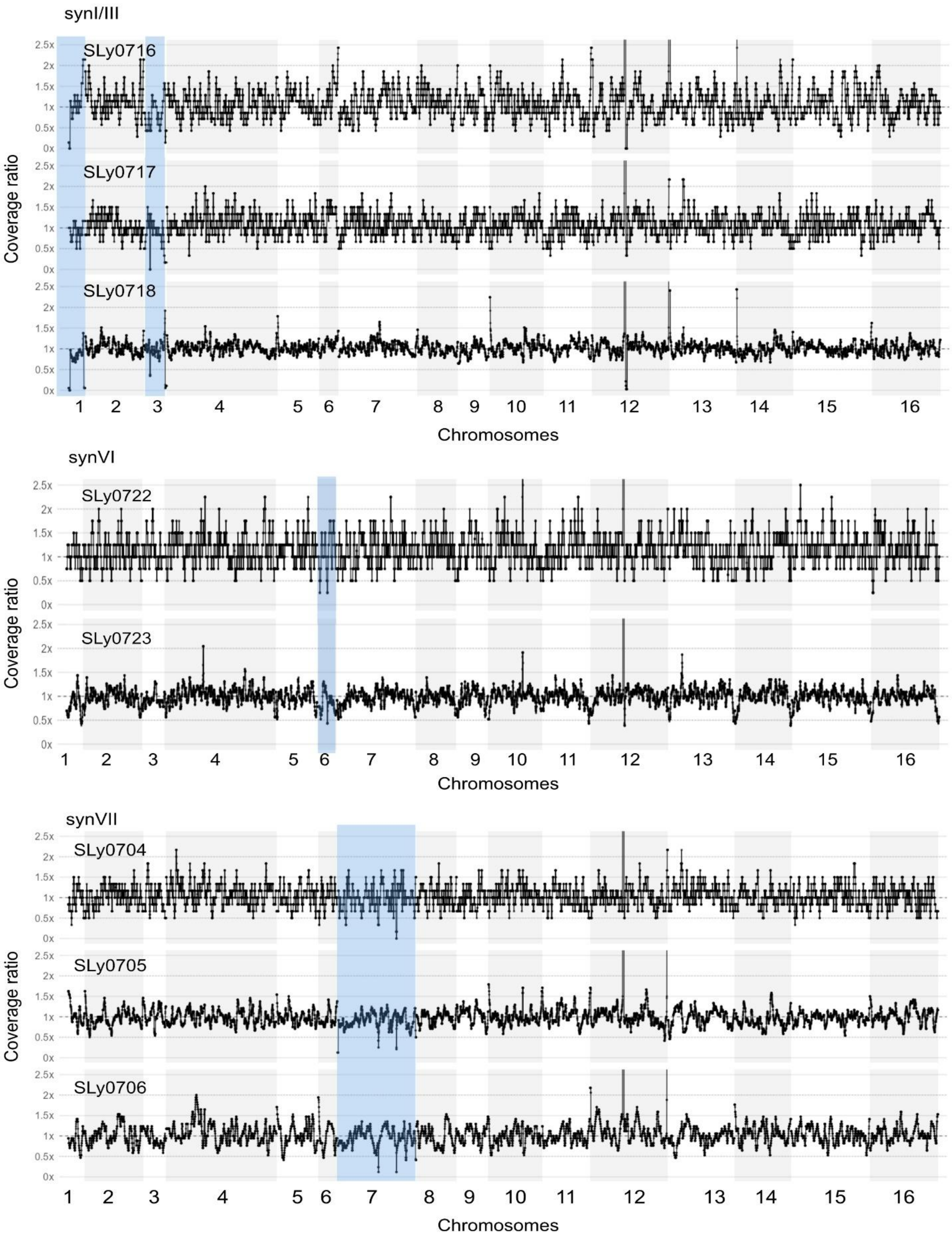


[Figure continous on following page]

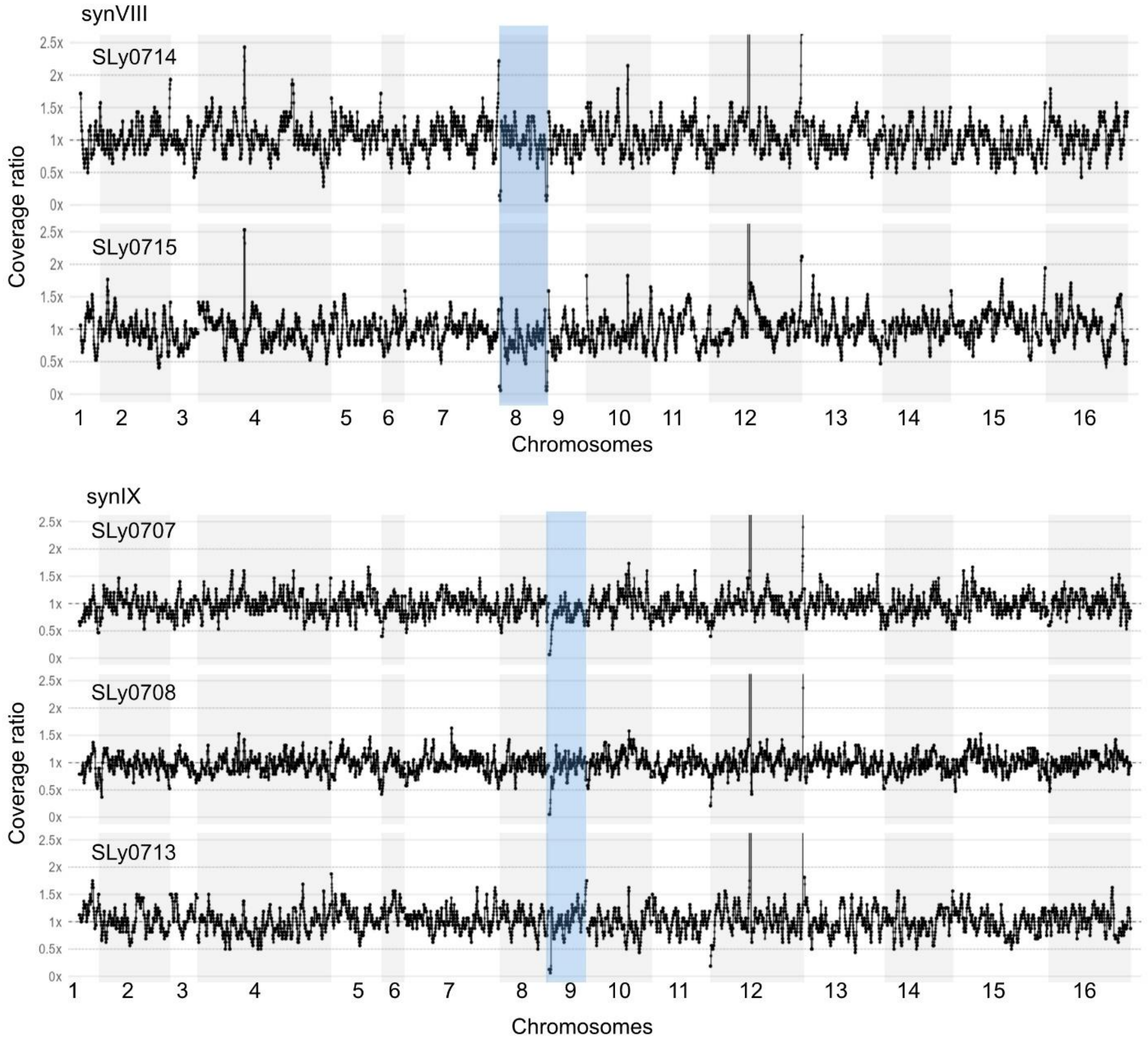


**Figure S12 | Analysis of aneuploidy of SCRaMbLed yeast synthetic strains.** Genome-wide sequencing coverage for each strain, binned in 1 kb non-overlapping windows. A ratio of 1.0 (dashed line) indicates coverage matching the own sample baseline, haploid level, deviations indicate copy-number gains or losses. Chromosomes are concatenated along the x-axis (alternating shading distinguishes chromosome boundaries) in the order indicated by numbers in the x-axis. Highlightedn blue synthetic yeast chromosomes.

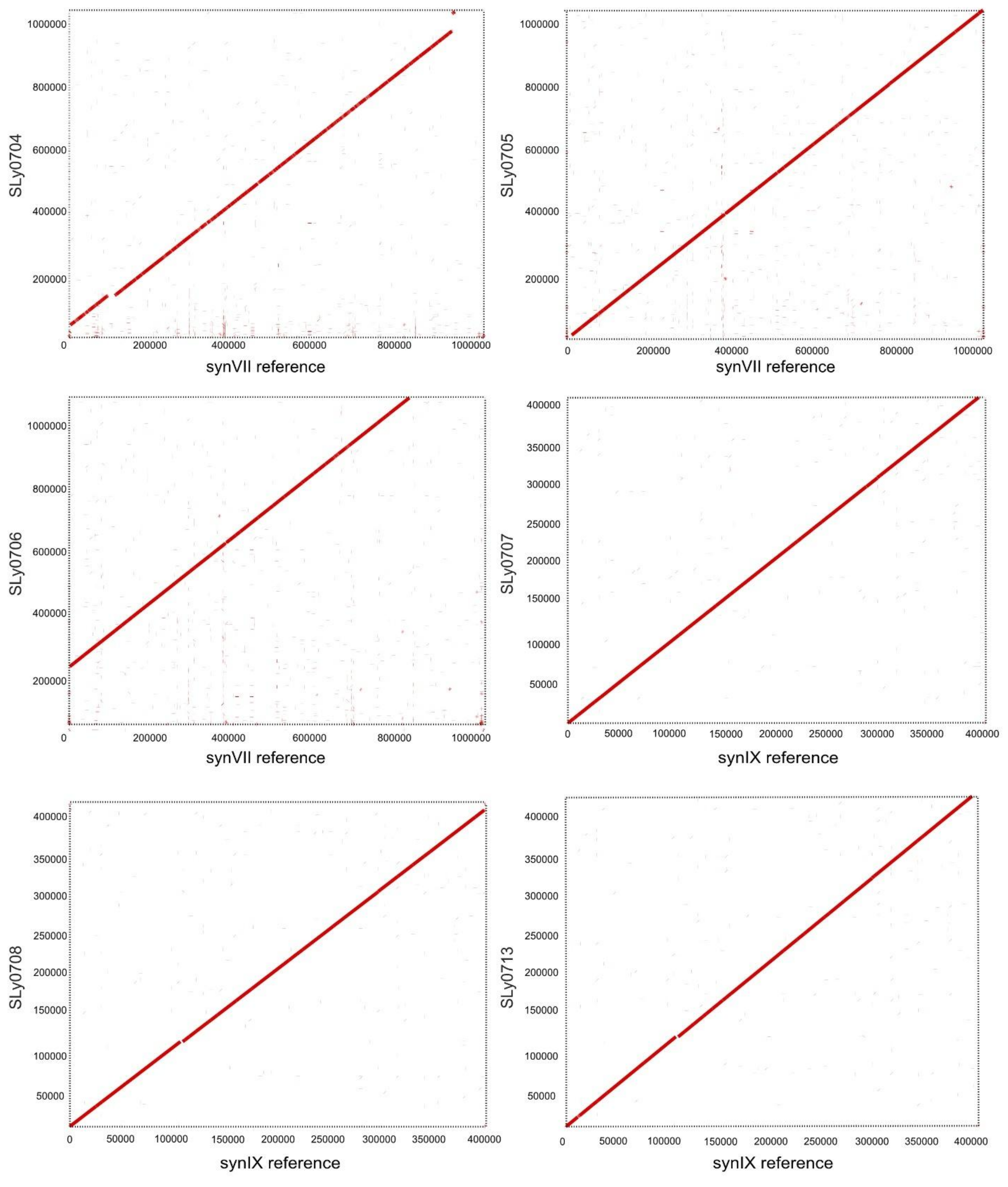


[Figure continous on following page]

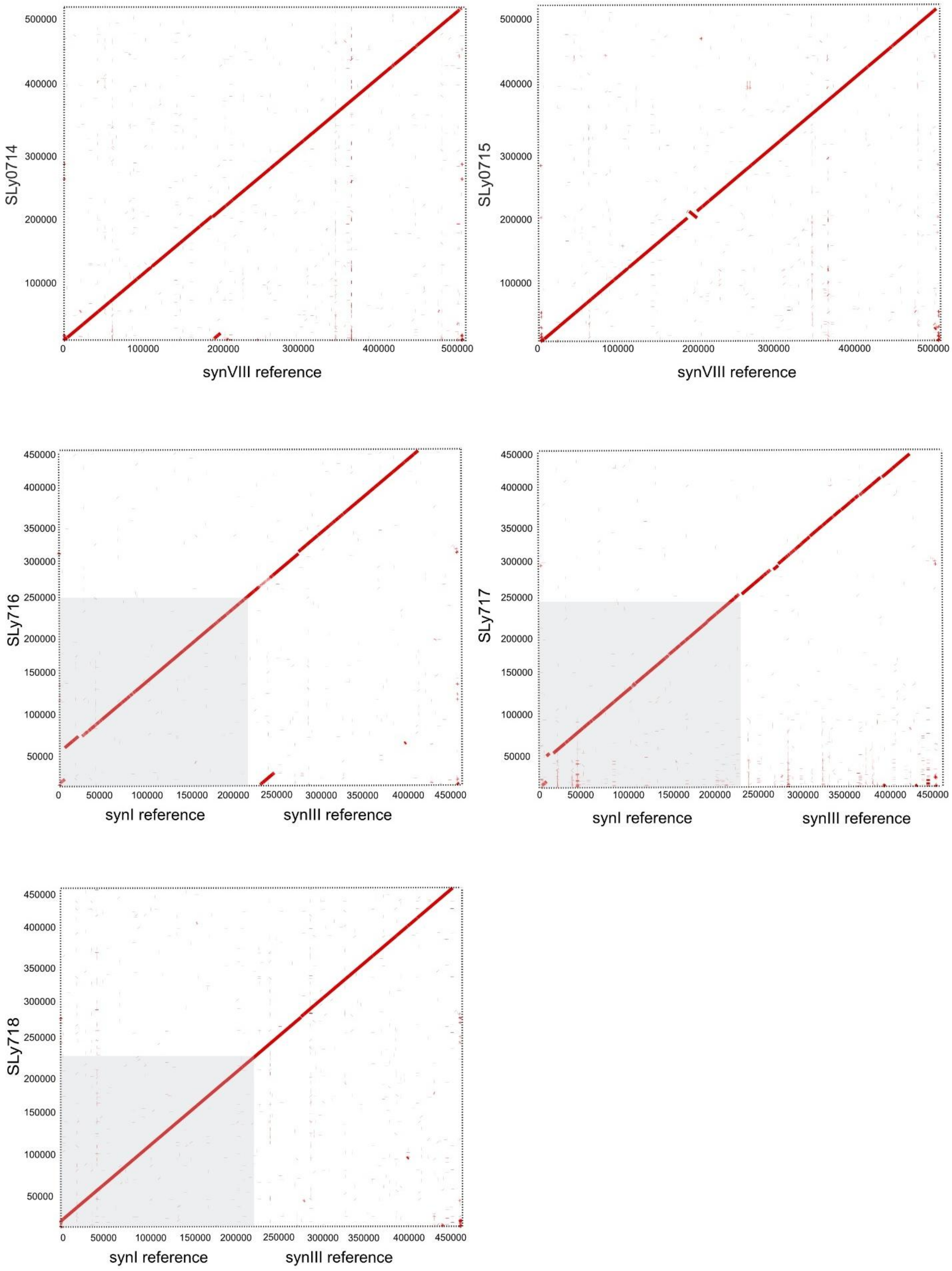


[Figure continous on following page]

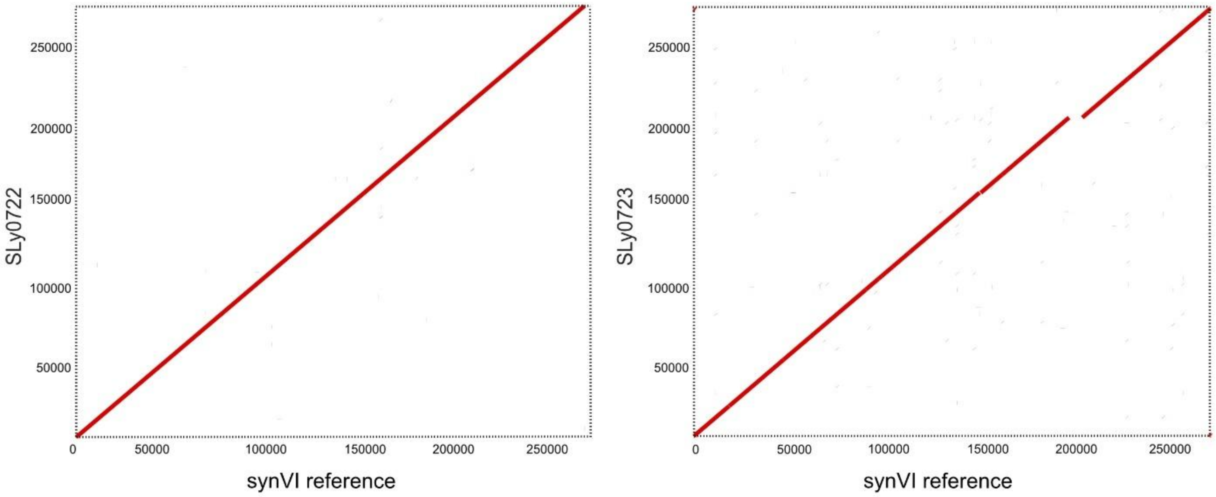


**Figure S13 | Dot plots showing structural rearrangements in SCRaMbLEd synthetic yeast strains.** Pairwise whole-chromosome alignments were generated using MUMmer between each SCRaMbLEd synthetic strain (y-axis) and its corresponding unscrambled reference. A continuous diagonal indicates collinear, unrearranged sequence, while breaks or discontinuities in the diagonal indicate structural rearrangements introduced by SCRaMbLE.

**Table S1 | Plasmids used in this study**

| ID | Type of plasmid | Propagation marker | Reference |
|---|---|---|---|
| **pSL0103** | Level 0 - CDS acceptor plasmid for N- & C-terminal tagging- | spectinomycin | [1] |
| **pSL0217** | Cre recombinase expression plasmid for SCRaMbLe experiments | ampicilin | Schindler Lab |
| **pSL0265** | Level 0 - Basic part Cyc1 terminator | spectinomycin | Schindler Lab |
| **pSL0836** | Level 0 - Basic part Gal10 promoter | spectinomycin | Schindler Lab |
| **pSL1005** | Level 0 - Basic part anchor protein CIS3 (N-terminal) | spectinomycin | This study |
| **pSL1006** | Level 0 - Basic part anchor protein AGA2 (N-terminal) | spectinomycin | This study |
| **pSL1007** | Level 0 - Basic part anchor protein CWP2 (N-terminal) | spectinomycin | This study |
| **pSL1008** | Level 0 - Basic part anchor protein TIP1 (N-terminal) | spectinomycin | This study |
| **pSL1009** | Level 0 - Basic part signal-peptide sequence α-pre-pro leader (based on [2]) | spectinomycin | This study |
| **pSL1010** | Level 0 - Basic part $sLac_{Amyco}$ (N-terminal) | spectinomycin | This study |
| **pSL1011** | Level 0 - Basic part anchor protein AGA2 (C-terminal) | spectinomycin | This study |
| **pSL1012** | Level 0 - Basic part anchor protein CWP2 (C-terminal) | spectinomycin | This study |
| **pSL1013** | Level 0 - Basic part anchor protein CIS3 (C-terminal) | spectinomycin | This study |
| **pSL1014** | Level 0 - Basic part anchor protein TIP1 (C-terminal) | spectinomycin | This study |
| **pSL1015** | Level 0 - Basic part $sLac_{Amyco}$ (C-terminal) | spectinomycin | This study |
| **pSL1017** | Level 1 - Acceptor vector | ampicillin | This study |
| **pSL1022** | Level 1 - Transcription unit $sLac_{Amyco}$-TIP1 (N-terminal) | ampicilin | This study |
| **pSL1023** | Level 1 - Transcription unit $sLac_{Amyco}$-AGA2(N-terminal) | ampicilin | This study |
| **pSL1024** | Level 1 - Transcription unit $sLac_{Amyco}$-CIS3 (N-terminal) | ampicilin | This study |
| **pSL1025** | Level 1 - Transcription unit $sLac_{Amyco}$-CWP2 (N-terminal) | ampicilin | This study |
| **pSL1030** | Level 1 - Transcription unit $sLac_{Amyco}$-TIP1 (C-terminal) | ampicilin | This study |
| **pSL1031** | Level 1 - Transcription unit $sLac_{Amyco}$-AGA2(C-terminal) | ampicilin | This study |
| **pSL1032** | Level 1 - Transcription unit $sLac_{Amyco}$-CIS3 (C-terminal) | ampicilin | This study |
| **pSL1033** | Level 1 - Transcription unit $sLac_{Amyco}$-CWP2 (C-terminal) | ampicilin | This study |
| **pSL1131** | Level 1 - Acceptor vector | ampicilin | This study |
| **pSL1133** | Level 0 - $sLac_{Sspir}$ (N-terminal) | spectinomycin | This study |
| **pSL1134** | Level 0 - CDS sLac *Halococcus sediminicola* (N- terminal) | spectinomycin | This study |
| **pSL1135** | Level 0 - CDS sLac *Methylomicrobium* sp016745425 (N- terminal) | spectinomycin | This study |
| **pSL1136** | Level 0 - CDS sLac *Actinoplanes xinjiangensis* (N- terminal) | spectinomycin | This study |
| **pSL1137** | Level 0 - CDS sLac *Planosporangium flavigriseum* (N- terminal) | spectinomycin | This study |
| **pSL1138** | Level 0 - CDS sLac *Micromonospora violae* (N- terminal) | spectinomycin | This study |
| **pSL1139** | Level 0 - CDS sLac SIRX01 sp004563715 (N- terminal) | spectinomycin | This study |
| **pSL1140** | Level 0 - CDS sLac *Pseudomonas resinovorans* (N- terminal) | spectinomycin | This study |
| **pSL1141** | Level 0 - CDS sLac *Actinacidiphila bryophytorum* (N- terminal) | spectinomycin | This study |
| **pSL1142** | Level 1 - $sLac_{Sspir}$ -AGA2 (N- terminal) | ampicilin | This study |
| **pSL1143** | Level 1 - Transcription unit sLac *H. sediminicola*-AGA2 (N-terminal) | ampicilin | This study |
| **pSL1144** | Level 1 - Transcription unit sLac *M.* sp016745425-AGA2 (N-terminal) | ampicilin | This study |

| ID | Type of plasmid | Propagation marker | Reference |
|---|---|---|---|
| **pSL1145** | Level 1 - Transcription unit sLac *A. xinjiangensis*-AGA2 (N-terminal) | ampicilin | This study |
| **pSL1146** | Level 1 - Transcription unit sLac *P. flavigriseum*-AGA2 (N-terminal) | ampicilin | This study |
| **pSL1147** | Level 1 - Transcription unit sLac *M. violae*-AGA2 (N-terminal) | ampicilin | This study |
| **pSL1148** | Level 1 - Transcription unit sLac SIRX01 sp004563715-AGA2 (N-terminal) | ampicilin | This study |
| **pSL1149** | Level 1 - Transcription unit sLac *P. resinovorans*-AGA2 (N-terminal) | ampicilin | This study |
| **pSL1150** | Level 1 - Transcription unit sLac *A. bryophytorum*-AGA2 (N-terminal) | ampicilin | This study |
| **pSL1151** | Level 1 - Transcription unit $sLac_{Sspir}$-CWP2 (N-terminal) | ampicilin | This study |
| **pSL1152** | Level 1 - Transcription unit sLac *H. sediminicola*-CWP2 (N-terminal) | ampicilin | This study |
| **pSL1153** | Level 1 - Transcription unit sLac *M.* sp016745425-CWP2 (N-terminal) | ampicilin | This study |
| **pSL1154** | Level 1 - Transcription unit sLac *A. xinjiangensis*-CWP2 (N-terminal) | ampicilin | This study |
| **pSL1155** | Level 1 - Transcription unit sLac *P. flavigriseum*-CWP2 (N-terminal) | ampicilin | This study |
| **pSL1156** | Level 1 - Transcription unit sLac *M. violae*-CWP2 (N-terminal) | ampicilin | This study |
| **pSL1157** | Level 1 - Transcription unit sLac SIRX01 sp004563715-CWP2 (N-terminal) | ampicilin | This study |
| **pSL1158** | Level 1 - Transcription unit sLac *P. resinovorans*-CWP2 (N-terminal) | ampicilin | This study |
| **pSL1159** | Level 1 - Transcription unit sLac *A. bryophytorum*-CWP2 (N-terminal) | ampicilin | This study |
| **pSL1160** | Level 1 - Transcription unit sLac $sLac_{Sspir}$-CIS3 (N-terminal) | ampicilin | This study |
| **pSL1161** | Level 1 - Transcription unit sLac *H. sediminicola*-CIS3 (N-terminal) | ampicilin | This study |
| **pSL1162** | Level 1 - Transcription unit sLac *M.* sp016745425-CIS3 (N-terminal) | ampicilin | This study |
| **pSL1163** | Level 1 - Transcription unit sLac *P. flavigriseum*-CIS3 (N-terminal) | ampicilin | This study |
| **pSL1164** | Level 1 - Transcription unit sLac *M. violae*-CIS3 (N-terminal) | ampicilin | This study |
| **pSL1165** | Level 1 - Transcription unit sLac SIRX01 sp004563715-CIS3 (N-terminal) | ampicilin | This study |
| **pSL1166** | Level 1 - Transcription unit sLac *P. resinovorans*-CIS3 (N-terminal) | ampicilin | This study |
| **pSL1167** | Level 1 - Transcription unit sLac *A. bryophytorum*-CIS3 (N-terminal) | ampicilin | This study |
| **pSL1168** | Level 1 - Transcription unit sLac *A. xinjiangensis*-CIS3 (N-terminal) | ampicilin | This study |
| **pSL1169** | Level 1 - Transcription unit $sLac_{Sspir}$-TIP1 (N-terminal) | ampicilin | This study |
| **pSL1170** | Level 1 - Transcription unit sLac *H. sediminicola*-TIP1 (N-terminal) | ampicilin | This study |
| **pSL1171** | Level 1 - Transcription unit sLac *M.* sp016745425-TIP1 (N-terminal) | ampicilin | This study |
| **pSL1172** | Level 1 - Transcription unit sLac *A. xinjiangensis*-TIP1 (N-terminal) | ampicilin | This study |
| **pSL1173** | Level 1 - Transcription unit sLac *P. flavigriseum*-TIP1 (N-terminal) | ampicilin | This study |
| **pSL1174** | Level 1 - Transcription unit sLac *M. violae*-TIP1 (N-terminal) | ampicilin | This study |
| **pSL1175** | Level 1 - Transcription unit sLac SIRX01 sp004563715-TIP1 (N-terminal) | ampicilin | This study |
| **pSL1176** | Level 1 - Transcription unit sLac *P. resinovorans*-TIP1 (N-terminal) | ampicilin | This study |
| **pSL1177** | Level 1 - Transcription unit sLac *A. bryophytorum*-TIP1 (N-terminal) | ampicilin | This study |
| **pSL1179** | Level 1 - Transcription unit sLac $sLac_{Amyco}$-CIS3 (N-terminal) evolved | ampicilin | This study |
| **pRSII413** | Control plasmid used for SCRaMbLe experiments | ampicilin | [3]<br>Addgene #35450 |
| **pRSII416** | Control plasmid used for SCRaMbLe experiments | ampicilin | [3]<br>Addgene #35456 |
| **pSCW11-CreEBD** | Cre recombinase expression plasmid for SCRaMbLe experiments | ampicilin | [4] |
| **pWS082** | Guide RNA entry plasmid | ampicilin | Addgene #90516 |

| ID | Type of plasmid | Propagation marker | Reference |
| --- | --- | --- | --- |
| **pWS158** | Cas9 expression plasmid | kanamycin | Addgene #90517 |

**Table S2 | Selected putative small bacterial laccases for experimental validation**

| Species | Ortho-group | Ref structure | TM-score (Chain_1) | TM-score (Chain_2) | LDDT | RMSD | Mol. weight (kDa) | Proline content (%) | Isolation source |
| --- | --- | --- | --- | --- | --- | --- | --- | --- | --- |
| *A. bryophytorum* | K0421 | PDB: 7Y8C | 0.8344 | 0.8507 | 0.62 | 1.90 | 52 | 6.68 | Endophytic |
| *H. sedimenticola* | K0421 | UniprotID: D4GPK6 | 0.9140 | 0.8221 | 0.78 | 2.41 | 67 | 7.82 | Marine sediment |
| *P. resinovorans* | K0421 | UniprotID: D4GPK6 | 0.8470 | 0.7833 | 0.56 | 2.33 | 63 | 8.15 | Soil |
| SIRX01 sp004563715 | K0421 | UniprotID: D4GPK6 | 0.9335 | 0.9304 | 0.57 | 2.21 | 61 | 7.35 | Hot spring sediment |
| *A. xinjiangensis* | K10535 | PDB: 3TA4 | 0.9415 | 0.7984 | 0.78 | 1.69 | 32 | 5.39 | Soil |
| *M.* sp016745425 | K10535 | PDB: 3TA4 | 0.8035 | 0.7228 | 0.67 | 2.17 | 32 | 5.42 | Rice rhizosphere |
| *M. violae* | K10535 | PDB: 3TA4 | 0.9362 | 0.7940 | 0.78 | 1.66 | 31 | 5.90 | Root of *Viola philippica* |
| *P. flavigriseum* | K10535 | PDB: 3TA4 | 0.9462 | 0.7920 | 0.77 | 1.24 | 33 | 4.38 | Evergreen broadleaved forest at Menghai |
| *S. spiralis* | K10535 | PDB: 3TA4 | 0.9376 | 0.7860 | 0.76 | 1.65 | 33 | 6.31 | Soil |

**Table S3 | Compounds detected by HPLC-MS after lignin degradation by $slac_{Sspir}$ and $slac_{Amyco}$**

| Compound number | Class | Putative identification name | Structure | Depolymerisation-related references |
|---|---|---|---|---|
| 1 | Benzaldehydes | P-formylphenol/4-hydroxy benzaldehyde | 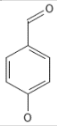 | [5-7] |
| 2 | Lignan | 4-[4-(4-hydroxy-3-methoxyphenyl)oxolan-3-yl]-2-methoxyphenol/lariciresinol | 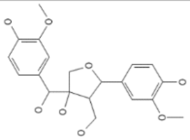 | NA |
| 3 | Aromatic ketone | 1-(4-hydroxyphenyl)propan-2-one | 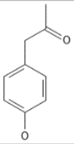 | NA |
| 4 | Benzaldehydes | Syringaldehyde | 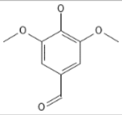 | [6, 8, 9] |
| 5 | Aromatic ketone | Disyringylketone | 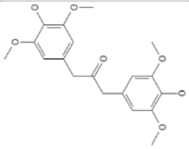 | [10] |
| 6 | Phenol | Vanillic acid | 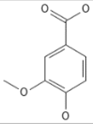 | [11] |
| 7 | Aromatic ketone | Gingerol | 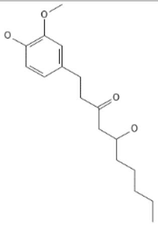 | NA |
| 8 | Lignan | Matairesinol | 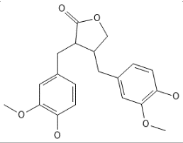 | NA |
| 9 | Benzaldehydes | 2-hydroxy-5-methoxybenzaldehyde | 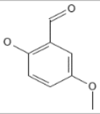 | NA |
| 10 | Aromatic oligomer | 4-{4-hydroxy-3-[(4-hydroxy-3-methoxyphenyl)methyl]but-1-en-1-yl}-2-methoxyphenol | 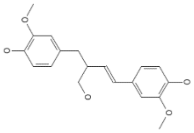 | NA |
| 11 | Aromatic oligomer | 4-{[5-(4-hydroxy-3-methoxyphenyl)-4-(hydroxymethyl)oxolan-3-yl]methyl}benzene-1,2-diol | 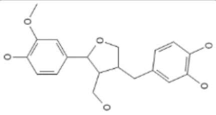 | NA |

| Compound number | Class | Putative identification name | Structure | Depolymerisation-related references |
|---|---|---|---|---|
| 12 | Aromatic oligomer | 4-{4-hydroxy-3-[(4-hydroxy-3-methoxyphenyl)methyl]-2-(hydroxymethyl)butyl}benzene-1,2-diol | 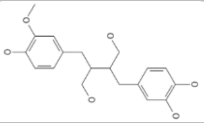 | NA |
| 13 | Aromatic oligomer | 3-hydroxy-5-methoxy-2,2-dimethyl-7-(2-phenylethyl)-3,4-dihydro-2H-1-benzopyran-8-carboxylic acid | 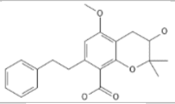 | NA |
| 14 | Aromatic oligomer | Pestalotether E | 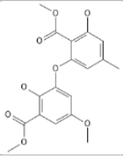 | NA |
| 15 | Aromatic oligomer (similar to stilbenoids) | 4-[(E)-2-(4-hydroxy-3,5-dimethoxyphenyl)ethenyl]-2,6-dimethoxyphenol | 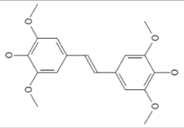 | NA |
| 16 | Phenol | Mequinol | 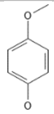 | NA |
| 17 | Aromatic ketone | Piceol | 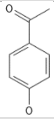 | NA |
| 18 | Phenol | Vanillactate | 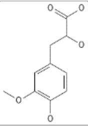 | NA |
| 19 | Lignan | Tanegool | 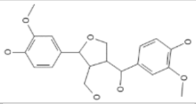 | NA |
| 20 | Lignan | Secoisolariciresinol | 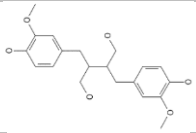 | [5] |
| 21 | Neolignan | 7R,8R-4,9,9'-trihydroxy-3,3'-Dimethoxy-8-O-4'-Neolignan | 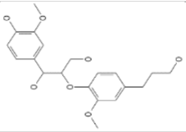 | NA |
| 22 | Benzaldehyde | Canillin | 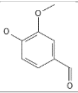 | **[5-7, 12, 13]** |
| 23 | Lignan | Pinoresinol | 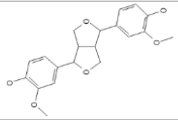 | NA |
| 24 | Benzoquinone | 6-[(2E,4E)-2,4,10-dodecatrienyl]-2,3-dimethoxy-5-methyl-1,4-benzoquinone | 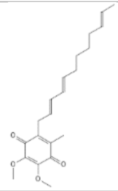 | NA |

| Compound number | Class | Putative identification name | Structure | Depolymerisation-related references |
|---|---|---|---|---|
| 25 | Aromatic oligomer | 4-({4-[(4-hydroxy-3-methoxyphenyl)methyl]oxolan-3-yl}methyl)-2-methoxyphenol | 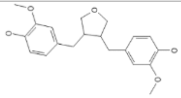 | NA |
| 26 | Aromatic oligomer | 4-[5-(4-hydroxy-3-methoxyphenyl)-3,4-dimethylfuran-2-yl]-2-methoxyphenol | 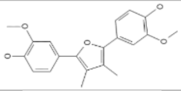 | NA |
| 27 | Aromatic oligomer | 4-[2-(4-hydroxy-3-methoxyphenyl)ethyl]-2-methoxyphenol | 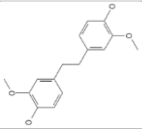 | NA |

**Table S4 | Yeast strains used in this study**

| ID | Genotype | Strain background | Parental strain | Purpose | Reference |
|---|---|---|---|---|---|
| BY4741 | MATa his3Δ1 leu2Δ0 met15Δ0 ura3Δ0 | S288c | S288c | Wild-type reference strain | [14] |
| BY4742 | lys2Δ0 ura3Δ0 his3Δ1 leu2Δ0 | S288c | S288c | Wild-type reference strain | [14] |
| yCTC002 (synI-synIII) | BY4742 synI-synIII ho::SYN.tS(CGA)C | BY4742 | BY4742 | Yeast strain with synI fused to synIII | [15] |
| YZY085 (synI/III) | BY4742 synI synIII ho::SYN.tS(CGA)C | BY4742 | BY4742 | Yeast strain with synI and synIII | [15] |
| YZY166 (synII) | BY4741 synII | BY4741 | BY4741 | Yeast strain with synII | [16] |
| YZY454 (synIII) | BY4742 synIII ho::SYN.tS(CGA)C | BY4742 | BY4742 | Yeast strain with synIII | [17] |
| yWZ703 (synIV) | BY4741 synIV | BY4741 | BY4741 | Yeast strain with synIV | [18] |
| yXZX846 (synV) | BY4741 synV | BY4741 | BY4741 | Yeast strain with synV | [19] |
| yLM953 (synVI) | BY4741 synVI | BY4741 | BY4741 | Yeast strain with synVI | [20] |
| YZY767 (synVII) | BY4741 synVII | BY4741 | BY4741 | Yeast strain with synVII | [21] |
| ySLL217 (synVIII) | BY4741 synVIII | BY4741 | BY4741 | Yeast strain with synVIII | [22] |
| yLHM1601 (synIX) | BY4742 synIX | BY4742 | BY4742 | Yeast strain with synIX | [23, 24] |
| yYW115 (synX) | BY4741 synX | BY4741 | BY4741 | Yeast strain with synX | [25] |
| YZY1203 (synXI) | MATa his3Δ1 leu2Δ0 lys2Δ0 ura3Δ0 synXI [pRS413-chrXI_tRNA] | BY4741 | BY4741 | Yeast strain with synXI | [26] |
| yWZ036 (synXII) | BY4742 YLR151C.HIS3.YLR152C | BY4742 | BY4742 | Yeast strain with synXII | [27] |
| Yzc024 (synXIII) | MATa his3Δ1 leu2Δ0 lys2Δ0 MET15 ura3Δ0 synXIII [1-883749] | BY4741 | BY4741 | Yeast strain with synXIII | [28] |
| synXIV.78 (synXIV) | BY4741 synXIV | BY4741 | BY4741 | Yeast strain with synXIV | [29] |
| YZY042 (synXV) | BY4741 WT.8thTag.IRA2 WT.OSW1 LoxΔ.CPA1-5' UTR | NA | NA | Yeast strain with synXV | [30] |
| YZY821 (synXVI) | BY4741 synXVI | BY4741 | BY4741 | Yeast strain with synXVI | [31] |
| SLy0475 | BY4741 ΔSED1 | BY4741 | BY4741 | SED1 deletion in BY4741 | This study |
| SLy0519 | SLy0475 pSL1024 | BY4741 | SLy0475 | Yeast with indicated sLac expression plasmid | This study |

| ID | Genotype | Strain background | Parental strain | Purpose | Reference |
|---|---|---|---|---|---|
| SLy0517 | SLy0475 pSL1022 | BY4741 | SLy0475 | Yeast with indicated sLac expression plasmid | This study |
| SLy0518 | SLy0475 pSL1023 | BY4741 | SLy0475 | Yeast with indicated sLac expression plasmid | This study |
| SLy0519 | SLy0475 pSL1024 | BY4741 | SLy0475 | Yeast with indicated sLac expression plasmid | This study |
| SLy0520 | SLy0475 pSL1025 | BY4741 | SLy0475 | Yeast with indicated sLac expression plasmid | This study |
| SLy0546 | SLy0475 pSL1017 | BY4741 | SLy0475 | Yeast with indicated sLac expression plasmid | This study |
| SLy0748 | SLy0475 pSL1142 | BY4741 | SLy0475 | Yeast with indicated sLac expression plasmid | This study |
| SLy0749 | SLy0475 pSL1143 | BY4741 | SLy0475 | Yeast with indicated sLac expression plasmid | This study |
| SLy0750 | SLy0475 pSL1144 | BY4741 | SLy0475 | Yeast with indicated sLac expression plasmid | This study |
| SLy0751 | SLy0475 pSL1145 | BY4741 | SLy0475 | Yeast with indicated sLac expression plasmid | This study |
| SLy0752 | SLy0475 pSL1146 | BY4741 | SLy0475 | Yeast with indicated sLac expression plasmid | This study |
| SLy0753 | SLy0475 pSL1147 | BY4741 | SLy0475 | Yeast with indicated sLac expression plasmid | This study |
| SLy0754 | SLy0475 pSL1148 | BY4741 | SLy0475 | Yeast with indicated sLac expression plasmid | This study |
| SLy0755 | SLy0475 pSL1149 | BY4741 | SLy0475 | Yeast with indicated sLac expression plasmid | This study |
| SLy0756 | SLy0475 pSL1150 | BY4741 | SLy0475 | Yeast with indicated sLac expression plasmid | This study |
| SLy0757 | SLy0475 pSL1151 | BY4741 | SLy0475 | Yeast with indicated sLac expression plasmid | This study |
| SLy0758 | SLy0475 pSL1152 | BY4741 | SLy0475 | Yeast with indicated sLac expression plasmid | This study |
| SLy0759 | SLy0475 pSL1153 | BY4741 | SLy0475 | Yeast with indicated sLac expression plasmid | This study |
| SLy0760 | SLy0475 pSL1154 | BY4741 | SLy0475 | Yeast with indicated sLac expression plasmid | This study |
| SLy0761 | SLy0475 pSL1155 | BY4741 | SLy0475 | Yeast with indicated sLac expression plasmid | This study |
| SLy0762 | SLy0475 pSL1156 | BY4741 | SLy0475 | Yeast with indicated sLac expression plasmid | This study |
| SLy0763 | SLy0475 pSL1157 | BY4741 | SLy0475 | Yeast with indicated sLac expression plasmid | This study |
| SLy0764 | SLy0475 pSL1158 | BY4741 | SLy0475 | Yeast with indicated sLac expression plasmid | This study |
| SLy0765 | SLy0475 pSL1159 | BY4741 | SLy0475 | Yeast with indicated sLac expression plasmid | This study |
| SLy0766 | SLy0475 pSL1160 | BY4741 | SLy0475 | Yeast with indicated sLac expression plasmid | This study |
| SLy0767 | SLy0475 pSL1161 | BY4741 | SLy0475 | Yeast with indicated sLac expression plasmid | This study |
| SLy0768 | SLy0475 pSL1162 | BY4741 | SLy0475 | Yeast with indicated sLac expression plasmid | This study |
| SLy0769 | SLy0475 pSL1163 | BY4741 | SLy0475 | Yeast with indicated sLac expression plasmid | This study |
| SLy0770 | SLy0475 pSL1164 | BY4741 | SLy0475 | Yeast with indicated sLac expression plasmid | This study |
| SLy0771 | SLy0475 pSL1165 | BY4741 | SLy0475 | Yeast with indicated sLac expression plasmid | This study |
| SLy0772 | SLy0475 pSL1166 | BY4741 | SLy0475 | Yeast with indicated sLac expression plasmid | This study |
| SLy0773 | SLy0475 pSL1167 | BY4741 | SLy0475 | Yeast with indicated sLac expression plasmid | This study |
| SLy0774 | SLy0475 pSL1168 | BY4741 | SLy0475 | Yeast with indicated sLac expression plasmid | This study |
| SLy0775 | SLy0475 pSL1169 | BY4741 | SLy0475 | Yeast with indicated sLac expression plasmid | This study |
| SLy0776 | SLy0475 pSL1170 | BY4741 | SLy0475 | Yeast with indicated sLac expression plasmid | This study |
| SLy0777 | SLy0475 pSL1171 | BY4741 | SLy0475 | Yeast with indicated sLac expression plasmid | This study |
| SLy0778 | SLy0475 pSL1172 | BY4741 | SLy0475 | Yeast with indicated sLac expression plasmid | This study |
| SLy0779 | SLy0475 pSL1173 | BY4741 | SLy0475 | Yeast with indicated sLac expression plasmid | This study |
| SLy0780 | SLy0475 pSL1174 | BY4741 | SLy0475 | Yeast with indicated sLac expression plasmid | This study |

| ID | Genotype | Strain background | Parental strain | Purpose | Reference |
|---|---|---|---|---|---|
| SLy0781 | SLy0475 pSL1175 | BY4741 | SLy0475 | Yeast with indicated sLac expression plasmid | This study |
| SLy0782 | SLy0475 pSL1176 | BY4741 | SLy0475 | Yeast with indicated sLac expression plasmid | This study |
| SLy0783 | SLy0475 pSL1177 | BY4741 | SLy0475 | Yeast with indicated sLac expression plasmid | This study |
| SLy0784 | SLy0475 pSL1179 | BY4741 | SLy0475 | Yeast with indicated sLac expression plasmid | This study |
| SLy0704 | YZY767 (synVII) pSL1160 | synVII | SLy0792 | SCRaMbLED yeast with sLac expression plasmid | This study |
| SLy0705 | YZY767 (synVII) pSL1160 | synVII | SLy0792 | SCRaMbLED yeast with sLac expression plasmid | This study |
| SLy0706 | YZY767 (synVII) pSL1160 | synVII | SLy0792 | SCRaMbLED yeast with sLac expression plasmid | This study |
| SLy0707 | yLHM1601 (synIX) pSL1160 | synIX | SLy0794 | SCRaMbLED yeast with sLac expression plasmid | This study |
| SLy0708 | yLHM1601 (synIX) pSL1160 | synIX | SLy0794 | SCRaMbLED yeast with sLac expression plasmid | This study |
| SLy0713 | yLHM1601 (synIX) pSL1160 | synIX | SLy0794 | SCRaMbLED yeast with sLac expression plasmid | This study |
| SLy0714 | ySLL217 (synVIII) pSL1160 | synVIII | SLy0793 | SCRaMbLED yeast with sLac expression plasmid | This study |
| SLy0715 | ySLL217 (synVIII) pSL1160 | synVIII | SLy0793 | SCRaMbLED yeast with sLac expression plasmid | This study |
| SLy0716 | YZY085 (synI/III) pSL1160 | synI/III | SLy0786 | SCRaMbLED yeast with sLac expression plasmid | This study |
| SLy0717 | YZY085 (synI/III) pSL1160 | synI/III | SLy0786 | SCRaMbLED yeast with sLac expression plasmid | This study |
| SLy0718 | YZY085 (synI/III) pSL1160 | synI/III | SLy0786 | SCRaMbLED yeast with sLac expression plasmid | This study |
| SLy0722 | yLM953 (synVI) pSL1160 | synVI | yLM953 | SCRaMbLED yeast with sLac expression plasmid | This study |
| SLy0723 | YZY767 (synVII) pSL1160 | synVII | SLy0792 | SCRaMbLED yeast with sLac expression plasmid | This study |
| SLy0785 | yCTC002 (synI-synIII) pSL0217 | synI-synIII | yCTC002 | Synthetic yeast with Cre expression plasmid | This study |
| SLy0786 | YZY085 (synI/III) pSL0217 | synI/III | YZY085 | Synthetic yeast with Cre expression plasmid | This study |
| SLy0787 | YZY166 (synII) pSL0217 | synII | YZY166 | Synthetic yeast with Cre expression plasmid | This study |
| SLy0788 | YZY454 (synIII) pSL0217 | synIII | YZY454 | Synthetic yeast with Cre expression plasmid | This study |
| SLy0789 | yWZ703 (synIV) pSL0217 | synIV | yWZ703 | Synthetic yeast with Cre expression plasmid | This study |
| SLy0790 | yXZX846 (synV) pSL0217 | synV | yXZX846 | Synthetic yeast with Cre expression plasmid | This study |
| SLy0791 | yLM953 (synVI) pSL0217 | synVI | yLM953 | Synthetic yeast with Cre expression plasmid | This study |
| SLy0792 | YZY767 (synVII) pSL0217 | synVII | YZY767 | Synthetic yeast with Cre expression plasmid | This study |
| SLy0793 | ySLL217 (synVIII) pSL0217 | synVIII | ySLL217 | Synthetic yeast with Cre expression plasmid | This study |
| SLy0794 | yLHM1601 (synIX) pSL0217 | synIX | yLHM1601 | Synthetic yeast with Cre expression plasmid | This study |
| SLy0795 | yYW115 (synX) pSL0217 | synX | yYW115 | Synthetic yeast with Cre expression plasmid | This study |
| SLy0796 | YZY1203 (synXI) pSCW11-CreEBD | synXI | YZY1203 | Synthetic yeast with Cre expression plasmid | This study |

| ID | Genotype | Strain background | Parental strain | Purpose | Reference |
|---|---|---|---|---|---|
| SLy0797 | yWZ036 (synXII) pSCW11-CreEBD | synXII | yWZ036 | Synthetic yeast with Cre expression plasmid | This study |
| SLy0798 | Yzc024 (synXIII) pSL0217 | synXIII | Yzc024 | Synthetic yeast with Cre expression plasmid | This study |
| SLy0799 | synXIV.78 (synXIV) pSCW11-CreEBD | synXIV | synXIV.78 | Synthetic yeast with Cre expression plasmid | This study |
| SLy0800 | YZY042 (synXV) pSL0217 | synXV | YZY042 | Synthetic yeast with Cre expression plasmid | This study |
| SLy0801 | YZY821 (synXVI) pSL0217 | synXVI | YZY821 | Synthetic yeast with Cre expression plasmid | This study |
| SLy0802 | yCTC002 (synI-synIII) pRSII413 | synI-synIII | yCTC002 | Synthetic yeast with pRS-based control plasmid | This study |
| SLy0803 | YZY085 (synI/III) pRSII413 | synI/III | YZY085 | Synthetic yeast with pRS-based control plasmid | This study |
| SLy0804 | YZY166 (synII) pRSII413 | synII | YZY166 | Synthetic yeast with pRS-based control plasmid | This study |
| SLy0805 | YZY454 (synIII) pRSII413 | synIII | YZY454 | Synthetic yeast with pRS-based control plasmid | This study |
| SLy0806 | yWZ703 (synIV) pRSII413 | synIV | yWZ703 | Synthetic yeast with pRS-based control plasmid | This study |
| SLy0807 | yXZX846 (synV) pRSII413 | synV | yXZX846 | Synthetic yeast with pRS-based control plasmid | This study |
| SLy0808 | yLM953 (synVI) pRSII413 | synVI | yLM953 | Synthetic yeast with pRS-based control plasmid | This study |
| SLy0809 | YZY767 (synVII) pRSII413 | synVII | YZY767 | Synthetic yeast with pRS-based control plasmid | This study |
| SLy0810 | ySLL217 (synVIII) pRSII413 | synVIII | ySLL217 | Synthetic yeast with pRS-based control plasmid | This study |
| SLy0811 | yLHM1601 (synIX) pRSII413 | synIX | yLHM1601 | Synthetic yeast with pRS-based control plasmid | This study |
| SLy0812 | yYW115 (synX) pRSII413 | synX | yYW115 | Synthetic yeast with pRS-based control plasmid | This study |
| SLy0813 | YZY1203 (synXI) pRSII416 | synXI | YZY1203 | Synthetic yeast with pRS-based control plasmid | This study |
| SLy0814 | yWZ036 (synXII) pRSII416 | synXII | yWZ036 | Synthetic yeast with pRS-based control plasmid | This study |
| SLy0815 | Yzc024 (synXIII) pRSII413 | synXIII | Yzc024 | Synthetic yeast with pRS-based control plasmid | This study |
| SLy0816 | synXIV.78 (synXIV) pRSII413 | synXIV | synXIV.78 | Synthetic yeast with pRS-based control plasmid | This study |
| SLy0817 | YZY042 (synXV) pRSII413 | synXV | YZY042 | Synthetic yeast with pRS-based control plasmid | This study |
| SLy0818 | YZY821 (synXVI) pRSII413 | synXVI | YZY821 | Synthetic yeast with pRS-based control plasmid | This study |

**Table S5 | Relevant oligonucleotides used in this study**

| ID | Sequence (5´→ 3´) | Description |
|---|---|---|
| **SLo5667** | TACGTCTCAGACTTTGAAGCTCCAGAATC CGACAACGGGTTTTGAGA | Forward primer for sgRNA cloning to target SED1 in BY4741 |
| **SLo5668** | GTACGTCTCAAAACCCGTTGTCGGATTCT GGAGCTTCAAAGTCTGAGA | Reverse primer for sgRNA cloning to target SED1 in BY4741 |
| **SLo5669** | TACGTCTCAGACTTTGATACGTTCTCTAT GGAGGAGTTTTGAGAC | Forward primer for sgRNA cloning to target SED1 in BY4741 |
| **SLo5670** | GTACGTCTCAAAACTCCTCCATAGAGAAC GTATCAAAGTCTGAGA | Reverse primer for sgRNA cloning to target SED1 in BY4741 |
| **SLo5671** | CTACCTTCCATACACCACTGATTGC | Forward primer for amplification of repair template by overlap extension PCR for SED1 deletion in BY4741 |
| **SLo5672** | GCGAACGTATTTTATTTTGCTTGTCTTTG | Reverse primer for amplification of repair template by overlap extension PCR for SED1 deletion in BY4741 |
| **SLo5673** | CAAAGACAAGCAAAATAAAATACGTTCGC CGGTGGTGTTTGACACATCCG | Forward primer for amplification of repair template by overlap extension PCR for SED1 deletion in BY4741 |
| **SLo5674** | TACTGGCGTTCTCCATTTTGCTTAG | Reverse primer for amplification of repair template by overlap extension PCR for SED1 deletion in BY4741 |
| **SLo5675** | CGAGAAAGCTTAGCCCCGAGG | Forward primer to verify the deletion of SED1 by colony PCR |
| **SLo5676** | GCACAGCTGGATCCTCATAGC | Reverse primer to verify the deletion of SED1 by colony PCR |
| **SLo5731** | GCCAGCATTGCTGCTAAAGAAG | Forward primer for ep-PCR of small laccases |
| **SLo5732** | CAGGAAACGCAACGGATATTGAGTC | Reverse primer for ep- PCR of small laccases |
| **SLo5748** | CCCAGCTTCAGCCTCTCTTTTC | Forward primer for plasmid amplification (N-terminal constructions) for homologous recombination in yeast (related to directed evolution) |
| **SLo5749** | CAGGAACTGACAACTATATGCGAGC | Reverse primer for plasmid amplification (N-terminal constructions) for homologous recombination in yeast (related to directed evolution) |
| **SLo5750** | GCAATCAACAAGGCTGACAGC | Forward primer for sgRNA cloning to target Δsed1 deletion in BY4741 |
| **SLo5978** | GATACTTCGGCTGCCGAAACTG | Reverse primer for sgRNA cloning to target Δsed1 deletion in BY4741 |
| **SLo5979** | GCTTCAGCCTCTCTTTTCTC | Forward primer for sgRNA cloning to target Δsed1 deletion in BY4741 |
| **SLo5980** | GGGAATTAATGTCCCCAATGATAGC | Reverse primer for sgRNA cloning to target Δsed1 deletion in BY4741 |
| **SLo5981** | GCTGTTGTGGTCGCTTGG | Forward primer for amplification of repair template by overlap extension PCR for SED1 deletion in BY4741 |
| **SLo5982** | GTACGGCGTCGATTCTAAAG | Reverse primer for amplification of repair template by overlap extension PCR for SED1 deletion in BY4741 |
| **SLo5983** | CTTGAGCGGTAGCTGCAGA | Forward primer for amplification of repair template by overlap extension PCR for SED1 deletion in BY4741 |
| **SLo5984** | CCTGTCAAAAGTATGCATAGTGC | Reverse primer for amplification of repair template by overlap extension PCR for SED1 deletion in BY4741 |

**Table S6 | PFAM IDs selected for lignin degradation and assimilation using MEBS**

| PFAM_ID | Function/Name | Exemplary gene name or protein name [if applicable] | Category number (MEBS) | Description |
|---|---|---|---|---|
| **PF02578** | Multi-copper polyphenol oxidoreductase laccase | Laccase | 1 | Multicopper oxidoreductase |
| **PF07731** | Cu-oxidase_2 | Laccase | 1 | Multicopper oxidoreductase |
| **PF07732** | Cu-oxidase_3 | Laccase | 1 | Multicopper oxidoreductase |
| **PF00394** | Multicopper oxidase | NA | 1 | Multicopper oxidoreductase |
| **PF11895** | Fungal peroxidase | NA | 2 | Peroxidase |
| **PF00141** | Peroxidase | NA | 2 | Peroxidase |
| **PF05199** | GMC_oxred_C | NA | 3 | AAO |
| **PF00732** | GMC_oxred_N | NA | 3 | AAO |
| **PF02913** | FAD-oxidase_C | NA | 3 | AAO |
| **PF01565** | FAD_binding_4 | NA | 3 | AAO |
| **PF03594** | Benzoate membrane transport protein | NA | 4 | Transport |
| **PF21105** | DyP dimeric alpha+beta barrel domain | NA | 5 | Dye-peroxidase |
| **PF20628** | Dyp-type peroxidase, C-terminal | NA | 5 | Dye-peroxidase |
| **PF04261** | Dye peroxidase | NA | 5 | Dye-peroxidase |
| **PF03573** | Transport | NA | 4 | Transport |
| **PF03922** | Transport | NA | 4 | Transport |
| **PF07690** | MFS | NA | 4 | Transport |
| **PF03241** | HpaB | 4-hydroxyphenylacetate 3-hydroxylase C terminal | 6 | H-lignin |
| **PF11794** | HpaB_N | 4-hydroxyphenylacetate 3-hydroxylase N terminal | 6 | H-lignin |
| **PF01613** | Flavin_Reduct | NA | 6 | H-lignin |
| **PF00501** | AMP-binding | Fcs | 7 | Lignin assimilation |
| **PF13193** | AMP-binding_C | Fcs | 7 | Lignin assimilation |
| **PF13353** | 4Fe-4S single cluster domain | PhdA | 7 | Lignin assimilation |
| **PF00037** | 4Fe-4S binding domain | PhdA | 7 | Lignin assimilation |
| **PF04055** | Radical SAM superfamily | PhdA | 7 | Lignin assimilation |
| **PF05870** | Phenolic acid decarboxylase | PAD | 7 | Lignin assimilation |
| **PF00106** | short chain dehydrogenase | PhdE/ligD | 6 | H-lignin |
| **PF00171** | Aldehyde dehydrogenase | FAD1/LigV | 7 | Lignin assimilation |
| **PF00501** | AMP-binding enzyme | FerA | 7 | Lignin assimilation |
| **PF00378** | Enoyl-CoA hydratase/isomerase | Hydroxycinnamoyl-CoA hydratase-lyase FerB/fchl | 7 | Lignin assimilation |
| **PF00903** | Glyoxalase/Bleomycin resistance protein/Dioxygenase superfamily | NA | 6 | H-lignin |

| PFAM_ID | Function/Name | Exemplary gene name or protein name [if applicable] | Category number (MEBS) | Description |
|---|---|---|---|---|
| **PF22632** | Dihydroxybiphenyl dioxygenase BphC D1 | This domain is found in dihydroxybiphenyl dioxygenase BphC and related proteins. | 6 | H-lignin |
| **PF01575** | MaoC like domain | CouM | 6 | H-lignin |
| **PF17885** | SMO *Pseudomonas* | SMO-StyA | 6 | H-lignin |
| **PF01613** | SMO *Pseudomonas* | NA | 6 | H-lignin |
| **PF01977** | 3-octaprenyl-4-hydroxybenzoate carboxy-lyase Rift-related domain | BsdC/AroY | 7 | Lignin assimilation |
| **PF20696** | 3-octaprenyl-4-hydroxybenzoate carboxy-lyase C-terminal domain | BsdC | 7 | Lignin assimilation |
| **PF20695** | 3-octaprenyl-4-hydroxybenzoate carboxy-lyase N-terminal domain | BsdC/AroY | 7 | Lignin assimilation |
| **PF00890** | FAD binding domain | Vinyl-Phenol-Reductase | 6 | H-lignin |
| **PF16884** | ADH_N_2 | Oxidoreductase [Pseudomonas aeruginosa PAO1 | 7 | Lignin assimilation |
| **PF00107** | PF00107 | Oxidoreductase [Pseudomonas aeruginosa PAO2 | 7 | Lignin assimilation |
| **PF00501** | AMP-binding | Fcs | 7 | Lignin assimilation |
| **PF13193** | AMP-binding_C | Fcs | 7 | Lignin assimilation |
| **PF01571** | Aminomethyltransferase folate-binding domain | LigM, DesA | 8 | G-lignin |
| **PF00355** | Rieske [2Fe-2S] domain | VanA/LigX | 8 | G-lignin |
| **PF19112** | Vanillate O-demethylase oxygenase C-terminal domain | VanA | 8 | G-lignin |
| **PF22290** | Dimethylamine monooxygenase subunit DmmA-like, N-terminal domain | NA | 8 | G-lignin |
| **PF00111** | 2Fe-2S iron-sulfur cluster binding domain | VanB/GcoB | 7 | Lignin assimilation |
| **PF00067** | Cytochrome P450 | GcoA | 7 | Lignin assimilation |
| **PF00970** | Oxidoreductase FAD-binding domain | GcoB | 7 | Lignin assimilation |
| **PF00175** | Oxidoreductase NAD-binding domain | GcoB | 7 | Lignin assimilation |
| **PF19112** | Vanillate O-demethylase oxygenase C-terminal domain | LigM | 9 | S-lignin |
| **PF00355** | Rieske [2Fe-2S] domain | LigM | 9 | S-lignin |
| **PF00108** | Thiolase, N-terminal domain | PcaF | 7 | Lignin assimilation |
| **PF00561** | alpha/beta hydrolase fold | PcaD | 7 | Lignin assimilation |
| **PF01144** | Coenzyme A transferase | PcaI/J | 7 | Lignin assimilation |
| **PF02803** | Thiolase, C-terminal domain | PcaF | 7 | Lignin assimilation |
| **PF00775** | Dioxygenase | CatA/PcaH/PcaG | 7 | Lignin assimilation |
| **PF02426** | Muconolactone delta-isomerase | CatC | 7 | Lignin assimilation |

| PFAM_ID | Function/Name | Exemplary gene name or protein name [if applicable] | Category number (MEBS) | Description |
|---|---|---|---|---|
| **PF02746** | Mandelate racemase / muconate lactonizing enzyme, N-terminal domain | CatB | 7 | Lignin assimilation |
| **PF04444** | Catechol dioxygenase N terminus | CatA | 7 | Lignin assimilation |
| **PF13378** | Enolase C-terminal domain-like | CatB | 7 | Lignin assimilation |
| **PF00206** | Lyase | PcaB | 7 | Lignin assimilation |
| **PF02627** | Carboxymuconolactone decarboxylase family | PcaC | 7 | Lignin assimilation |
| **PF10397** | Adenylosuccinate lyase C-terminus | PcaH | 7 | Lignin assimilation |
| **PF12391** | Protocatechuate 3,4-dioxygenase beta subunit N terminal | PcaB | 7 | Lignin assimilation |
| **PF13380** | CoA binding domain | FerA, Fcs | 7 | Lignin assimilation |
| **PF13607** | Succinyl-CoA ligase like flavodoxin domain | FerA | 7 | Lignin assimilation |
| **PF00043** | Glutathione S-transferase, C-terminal domain | LigF, beta-esterases | 7 | Lignin assimilation |
| **PF02798** | Glutathione S-transferase, N-terminal domain | LigF, beta-esterases | 7 | Lignin assimilation |
| **PF08669** | Glycine cleavage T-protein C-terminal barrel domain | LigM | 7 | Lignin assimilation |
| **PF13417** | Glutathione S-transferase, N-terminal domain | LigG, beta-esterases | 7 | Lignin assimilation |
| **PF02900** | Catalytic LigB subunit of aromatic ring-opening dioxygenase | LigZ/LigB | 7 | Lignin assimilation |
| **PF04909** | Amidohydrolase | LigW | 7 | Lignin assimilation |
| **PF01408** | Oxidoreductase family, NAD-binding Rossmann fold | LigC | 7 | Lignin assimilation |
| **PF03737** | Aldolase/RraA | LigK | 7 | Lignin assimilation |
| **PF07746** | Aromatic-ring-opening dioxygenase LigAB, LigA subunit | LigA | 7 | Lignin assimilation |
| **PF13607** | Succinyl-CoA ligase like flavodoxin domain | Fcs | 7 | Lignin assimilation |
| **PF02900** | Catalytic LigB subunit of aromatic ring-opening dioxygenase | LigB | 7 | Lignin assimilation |
